# Single-Nucleus Neuronal Transcriptional Profiling of Male *C. elegans* Uncovers Regulators of Sex-Specific and Sex-Shared Behaviors

**DOI:** 10.1101/2024.12.12.628226

**Authors:** Katherine S. Morillo, Jonathan St. Ange, Yifei Weng, Rachel Kaletsky, Coleen T. Murphy

## Abstract

Sexual differentiation of the nervous system drives profound neurobiological and behavioral differences between the sexes across various organisms, including *Caenorhabditis elegans*. Using single-nucleus RNA sequencing, we profiled and compared adult male and hermaphrodite *C. elegans* neurons, generating an atlas of adult male-specific and sex-shared neurons. We expanded the molecular map of male-specific neurons, and identified highly dimorphic expression of GPCRs, neuropeptides, and ion channels. Our data demonstrate sex-shared neurons exhibit substantial heterogeneity between the sexes, while sex-specific neurons repurpose conserved molecular pathways to regulate dimorphic behaviors. We show that the PHD neurons display remarkable similarity to sex-shared AWA neurons, suggesting partial repurposing of conserved pathways, and that they and the GPCR SRT-18 may play a role in pheromone sensing. We further demonstrate that the ubiquitously expressed MAPK phosphatase *vhp-1* regulates both sex-specific and sex-shared behaviors. Our data provide a rich resource for discovering sex-specific transcriptomic differences and the molecular basis of sex-specific behaviors.

## Introduction

Across sexually reproducing species, including flies, mice, and humans^1–3^, biological sex impacts neuronal and cognitive function at multiple levels. Sexually dimorphic differences in cell number, gene expression, neuronal connectivity, and synaptic signaling result in differences in behavior and disease progression between the sexes. Yet most studies centered on neuronal biology and function using model organisms typically focus on one sex^4,5^ or on early developmental stages before the onset of many of these dimorphisms upon sexual maturation^6^.

*Caenorhabditis elegans* has two sexes, hermaphrodites (XX) and males (XO)^7^. The cell lineages of both sexes are invariant, and their neuronal connectomes have been fully mapped^8^. Hermaphrodites have 8 sex-specific neurons, while males have 93 sex-specific neurons, many of which are involved in male copulation behaviors^9^. The two sexes share 294 neurons, but many of these shared neurons still exhibit sexually dimorphic structural and functional features, such as differences in gene expression^10^, synaptic connectivity^7^ ^8^, and behavior (foraging, pheromone sensing, learning, and memory)^11–14^. Many of these behavioral differences are mediated by sex-specific regulation in a small subset of neurons^15–18^. Thus, identifying transcriptomic changes at the single-cell level is crucial to characterize these sexually dimorphic behaviors.

Bulk RNA-sequencing has been used to characterize the neuronal transcriptomes of both sexes throughout early development^19^ and in young and aged male neurons^20^, revealing extensive transcriptional variation between the sexes. However, while single-neuron transcriptional information exists for the L4 larval^6^ and adult hermaphrodite stages^21^, the adult male single-neuron transcriptome remains undescribed. We set out to fill this gap by performing single-nucleus RNA-sequencing to describe the transcriptome of young adult male neurons. In addition to assessing the male neuronal transcriptome, we also surveyed young adult hermaphrodite neuronal transcriptomes to identify sex-based differences and uncover potential behavioral regulators.

## Results

### High-throughput isolation of *C. elegans* males and hermaphrodites

In self-fertilizing populations, only ∼0.1% of *C. elegans* are males^22^; therefore, to obtain the large number of males required for single-nucleus RNA sequencing (snSeq), we crossed pan-neuronal histone-GFP-labeled hermaphrodites^21^ into a *him-8* mutant background, which produces ∼40% males^23^. Fluorescent labeling of neuronal nuclei was observed in both sexes (Figure 1A, S1), including in the male tail, where 69/93 male-specific neurons are located ^24,25^.

**Figure 1:**
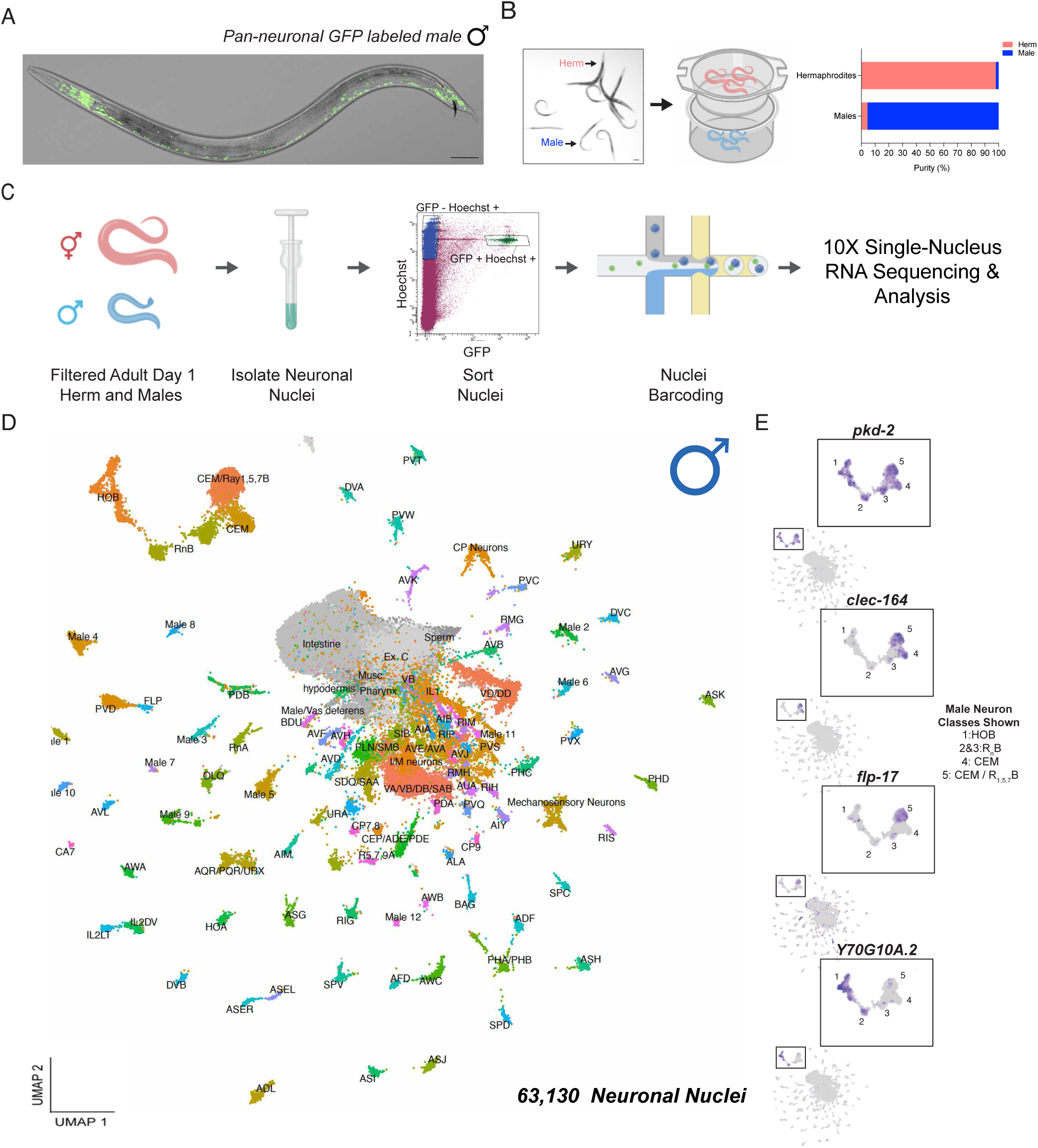
Single nucleus RNA sequencing of adult male *C. elegans* neurons. **(A)** Representative image of pan-neuronal GFP tagged *him-8* males (*Prgef-1::his-58::GFP; him-8)*. Scale bar: 50 µm **(B)** Representative image of synchronized day 1 adult hermaphrodites and males. 6-well tissue culture inserts with modified filters (35 µm) were used to separate hermaphrodites and males, obtaining isolated populations that were on average 95% enriched for each sex. Scale bar: 100 µm **(C)** Overview of nuclei isolation and sequencing. Pan-neuronal histone GFP-tagged *him-8* males and hermaphrodites were separated, nuclei were isolated, FACS sorted for Hoescht and DAPI positive nuclei, followed by barcoding, cDNA amplification, and library preparation using 10X genomics chromium X, then Illumina RNA sequenced. **(D)** UMAP of 63,130 male-specific neurons that passed quality control. **(E)** Feature plot of known markers expressed in RnB, CEM, and HOB neurons. *pkd-2* (expressed in all), *clec-164* (CEM- enriched), *flp-17* (CEM and R5,7,9B-enriched), and *Y70G10A.2* (HOB-enriched).

Day 1 of adulthood marks the onset of sexual maturity in *C. elegans;* both males and females start responding to mating cues at this stage^26^. Additionally, some male-specific neurons, such as the MCM and PHD neurons, arise from non-neuronal lineages upon sexual maturation^27,28^, thus would not be captured using pan-neuronal markers at earlier stages. Additionally, Day 1 of adulthood is preferred over L4, as hermaphrodites are unable to learn using a positive associative memory training paradigm prior to adulthood^21^. Therefore, we selected Day 1 adults as the ideal baseline to study both species-specific neurobiology and behavior, as well as sexual dimorphism in conserved behaviors, such as learning and memory. Using custom size-exclusion filters (Methods) to separate Day 1 adult males from Day 1 adult hermaphrodites, which are twice the size of males and too wide to pass through the filters^29^, we isolated ∼95% pure populations of each sex (Figure 1B, S1B).

### Neuronal single-nucleus RNA sequencing of day 1 adult males

Single-nucleus RNA sequencing is preferential to single-cell sequencing for neurons because the method reduces biases caused by dissociation problems and limits the loss of highly relevant, spatially-restricted RNAs in traditional cell isolation protocols by preserving nuclear transcripts^21^. Here, we applied our recently-developed method^21^ to isolate and sequence neuronal nuclei from Day 1 adult males, processed in parallel with age- and genotype-matched hermaphrodites. Nuclei were extracted by mechanical and chemical lysis, Hoechst-stained, and FACS-sorted based on dual GFP and Hoechst signals (Figure 1C, Methods). RNA was isolated, sequenced, then processed with Cell Ranger; SoupX-corrected^30^ ambient RNA was removed, and quality control metrics were assessed (Figure S1C-G). Given that the single-cell neuronal transcriptome of adult male *C. elegans* was previously uncharacterized, we first focused on systematic profiling of this male dataset. Across four male biological replicates, 63,130 male neuronal nuclei passed quality control filtering (Methods), and we obtained a median of 401 UMIs and 298 genes detected per nucleus (Figure S2).

### Cluster annotation and validation of male neuronal transcriptomes

*C. elegans* males have 387 neurons, of which 294 are shared between the two sexes and are grouped into 116 neuron classes based on morphological and connectivity similarities^31^, while 93 male-specific neurons fall into 27 distinct classes^32^. To assign neuronal identities to 110 clusters derived from our dataset, we employed two complementary statistical approaches, incorporating previously-identified neuronal markers and new markers identified in our recent hermaphrodite snSeq dataset^21^ to resolve neuronal identities (Methods). Further manual curation was used to confirm annotations and to resolve some neuron identities that were not readily apparent from statistical approaches. Using this strategy, we were able to annotate 74% of the clusters (Figure 1D, Table S1). Because the annotation process was primarily informed by hermaphrodite data, and 24% (93/387) of neurons in our dataset are expected to be male-specific, we posited that most of the remaining unannotated clusters represent male-specific neurons that are transcriptionally and functionally understudied. A smaller subset may represent highly dimorphic sex-shared neurons.

To assess the validity of our dataset, we examined a small subset of male-specific neurons that have been extensively studied over the past two decades — namely, the extracellular vesicle-releasing CEM, HOB, and ray B-type (RnB) neurons (Figure 1D,E). These neurons express the only two polycystins in *C. elegans, pkd-2* and *lov-1*, which are required for male mating behaviors, along with other functionally-relevant targets that now serve as hallmark markers for these neurons^33,34^. As expected, we observed robust expression of *pkd-2* across CEM, HOB, and RnB neurons, as well as enrichment of neuropeptide *flp-17* specifically in ray neurons R1B, R5B, R7B, as previously described^33^ (Figure 1E). We also detected high expression of the C-type lectin *clec-164,* a regulator of male sex drive^33^, in both CEM and RnB neurons. We also confirmed the predicted C-type lectin, Y70G10A.2, is highly enriched in HOB neurons (Figure 1E). These observations confirmed that our dataset faithfully recapitulates known male neuronal biology, validating both the quality of nuclei isolation and our cluster annotation approach. This foundation enabled the annotation of previously poorly-characterized male-specific neurons.

### Resolving identities of previously-uncharacterized male neurons

To annotate predicted male-specific neurons, we used a combination of transcriptional profiling, assessment of neuronal positional information upon promoter::GFP reporter generation, and manual curation of gene expression patterns. For example, the uncharacterized serpentine receptor *srd-66* was solely expressed in one of the unannotated clusters (cluster 105) (Figure 2A). To identify this cluster, we examined the expression pattern of *Psrd-66::GFP* animals; fluorescence was observed specifically in a neuron within the male ventral cord that is positionally consistent with the male-specific motor neuron CA7 (Figure 2B, left), with no detectable expression in hermaphrodites (Figure 2B, right). Furthermore, the glutamate transporter *eat-4,* which was previously shown to be expressed in CA7^35^, was also enriched in this cluster (Table S2), while *unc-47*, a GABAergic marker for adjacent CA/P6, CP7, and CA/P8 neurons but not CA7, is not expressed in this cluster (Table S2). Therefore, combining transcriptional information, neurotransmitter identity, and reporter validation, we are confident in our annotation of this cluster as male CA7 (Figure 2A, B).

**Figure 2:**
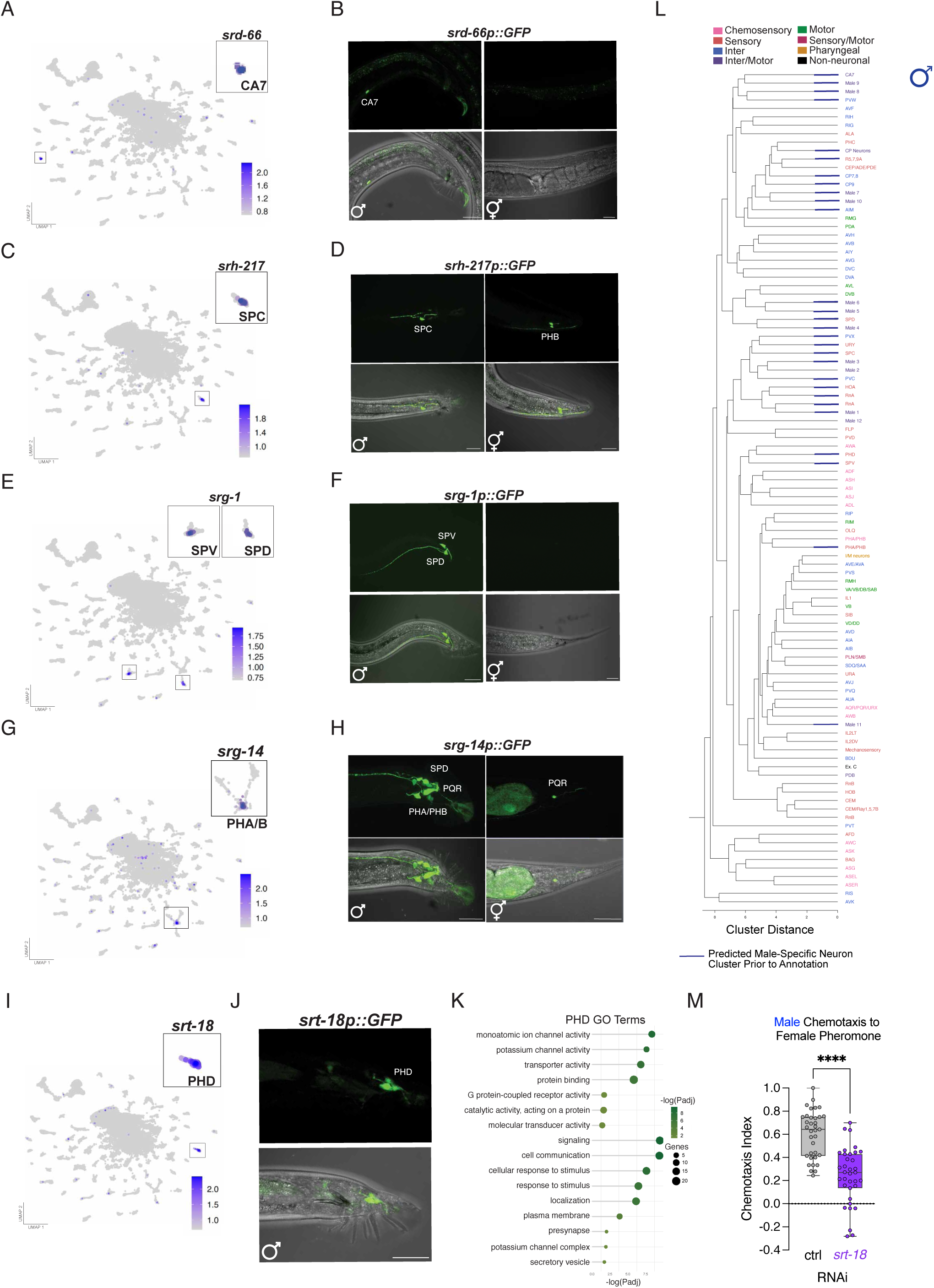
Annotation of previously poorly-characterized male-specific neurons. **(A)** Feature plot of *srd-66* expression. **(B)** Imaging of *srd-66p::GFP* shows corresponding expression and morphology for male CA7 neurons in Day 1 adult males; expression is male-specific, as no signal was detected in age-matched hermaphrodites. **(C)** Feature plot of *srh-217* expression. **(D)** Imaging of *srh-217p::GFP* shows corresponding expression and morphology for male SPC neurons, and hermaphrodite-specific PHB expression as previously reported. **(E)** Feature plot of *srg-1* expression. **(F)** Imaging of *srg-1p::GFP* shows corresponding expression and morphology for male-specific SPV and SPD neurons, and no expression in the hermaphrodite tail. **(G)** Feature plot of s*rg-14* expression. (**H**) Imaging of *srg-14p::GFP* shows corresponding expression and neuronal morphology in hermaphrodite PQR in the tail, while we observe at least 6 cell bodies expressing reporter in the male tail corresponding to SPD, PQR, and PHA/PHB neurons. **(I)** Feature plot of *srt-18* expression. **(J)** Imaging of *srt-18p::GFP* shows corresponding neuronal position and morphology for male PHD neurons. We did not observe any signal in hermaphrodites (not shown). **(K)** Significant Gene Ontology (GO) terms for PHD- enriched genes (Bonferroni padj < 0.05). Unpaired, two-tailed Student’s t-test. (**A, C, E, G, I**) Scale bar = SCT normalized expression level. (**B, D, F, H, J**) Scale bar = 25 µm. (**L)** Hierarchical dendrogram based on gene expression in each day 1 adult male neuron. Neurons are color-coded by functional subtype. Blue bars on branches indicate predicted male-specific/bias neurons. **(M)** Males’ ability to chemotaxis to female pheromone is attenuated upon *srt-18* neuron-specific RNAi knockdown. Chemotaxis assay performed using males from TU3595 neuron-specific RNAi-sensitive strain. N = 4 biological replicates. 5-10 chemotaxis plates per replicate, ∼30-50 worms per plate. *p<0.0001.

We also identified all three male spicule neurons - SPC, SPV and SPD - as distinct transcriptional clusters (clusters 60, 54, and 74, respectively) (Figure 1D, Table S1). These ciliated neurons innervate the male spicules and regulate vulva locating behaviors^36^. Consistent with their sensory identity, we observe high expression of ciliated neuron markers across all three clusters (Table S3), including *osm-6,* which was previously detected in spicule neurons^37^. Within these clusters, the serpentine receptor *srh-217* was highly enriched (∼40-fold over other neurons) in the cluster (number 60) corresponding to SPC (Figure 2C, D, Table S2). To validate this assignment, we generated *Psrh-217::*GFP, and observed fluorescence in neurons with distinct dendrites terminating near the spicule shaft (Figure 2D), consistent with SPC morphology^38^. *srh-217* was also expressed in the ASJ and AIB in both sexes (Figure S4A), which was previously shown only in hermaphrodites^6^. In the tail, *srh-217* was expressed in hermaphrodite PHB neurons, as previously reported, but not in male PHB neurons, consistent with our transcriptional data^6,21^ (Figure 2D, Table S2).

The serpentine receptor *srg-1* was highly enriched in the male SPD and SPV neurons, both of which exhibit clear processes running down the length of the spicule shaft (Figure 2E, F). We also detected expression of *srg-1* in the ASK chemosensory neurons in both sexes, as previously reported in hermaphrodites^6,21^. Additionally, SPV and SPD, but not SPC neurons, expressed the serpentine receptor *sra-1*, providing further distinction^39^. SPD neurons were distinguished from SPV by the enrichment of neuropeptide *flp-3* in SPV, as previously observed^40^. Together, these transcriptional and anatomical features validate our annotations of all three spicule neurons.

One unannotated male-biased cluster (number 91) was highly enriched for ciliated neuron markers and the serpentine receptor *srg-14* (Figure 2G, Table S3), which was previously detected in an unidentified male neuron pair and a few other neurons including, URX, AQR, PQR, and ASJ^39^ (Table S2). Interestingly, our statistical test assigned this cluster to the sex-shared sensory PHA/PHB neurons using L4-hermaphrodite markers (Table S1) but observed lack of overlap with adult hermaphrodite-based markers, suggesting extensive sexual dimorphism in adults. *Psrg-14::GFP* was highly expressed in at least 6 cell bodies in the male tail (Figure 2H), consistent with expression in PHA/PHB, SPD, and PQR tail neurons observed in our snSeq data. In the hermaphrodite tail, we only observed expression in PQR (Figure 2H, S2B). These findings support a model in which PHA and/or PHB neurons become highly dimorphic, and contribute to male-specific behaviors regulated by these neurons, such as mate-searching^32,33^.

The PHD neurons, a recently-discovered male ciliated neuron pair that arise from glial transdifferentiation during sexual maturation^43^, were also identified among our unannotated clusters. The chemoreceptor *srt-18* was highly enriched in cluster 34 and expressed 200-fold lower in cluster 96 (identified as AWB neurons) (Figure 2I, Table S2). *Psrt-18::GFP* showed clear expression of neurons in the lumbar ganglia and morphology, suggestive of the male PHD neurons^43^ (Figure 2J), identifying cluster 34 as the male PHD neurons. The PHD exhibited strong expression of ciliated neuron markers (*dyf* and *osm* genes; Table S3), the immunoglobulin domain-containing protein *oig-8,* and dense-core vesicle secretion factor *ida-1*^28,44^ (Table S2). Additionally, we detected >45 neuropeptide genes, including the highest-expressed neuropeptide, *nlp-50*, along with the vesicular acetylcholine transporter *unc-17,* consistent with peptidergic and cholinergic activity of the PHD neurons. Gene ontology analysis showed genes related to ion channel activity, signal transduction, response to stimulus, and localization (Figure 2K). Together, the data suggest that the PHD neurons are ciliated, cholinergic, and possibly chemosensory neurons, providing a molecular characterization of these recently-discovered male-specific class.

### Expansion of male-specific neuronal atlas

*C. elegans* males have 52 sex-specific ciliated sensory neurons^45^. Among these are the hook neuron HOB, along with the 4 CEM, 18 RnB, 2 PHD, and 3 spicule-associated SPC/SPV/SPD neurons (6), all which are annotated in our dataset. The remaining ciliated sensory neurons include the 18 ray A-type neurons, the hook neuron HOA, and one of three pairs of postcloacal sensilla (p.c.s) neurons, PCA ^46^. All these neuron types regulate male mating behaviors, such as vulva prodding, spicule insertion, and response to mates^36,46^. The RnA neurons share a common structure and function to their adjacent RnB counterparts, and are housed in the same acellular cuticular fan, but are morphologically^38^ and transcriptionally^33,47^ distinct, notably lacking expression of genes such as *lov-1* and *pkd-2* ^33^.

Based on these distinctions, we hypothesized that the remaining unannotated, ciliated gene enriched clusters in our dataset most likely correspond to RnA, HOA, and PCA neurons (Table S3). Supporting this notion, several of these clusters (numbers 29 and 41) were highly enriched in ciliary tubulin*, tbb-4* and *tba-9,* previously shown to be expressed in RnA but not RnB neurons^48^, consistent with our data (Table S2). Moreover, high expression of *mab-21,* a regulator of sensory ray differentiation expressed in all A-type rays^49^ and HOA, was also observed.

One cluster (number 102) showed enrichment for regulators of dopamine biosynthesis, dopamine transporter *dat-1* and tyrosine hydroxylase *cat-2,* consistent with neurotransmitter identity of the only male-specific dopaminergic neurons rays 5A, 7A, and 9A ^35,50^. Another cluster (number 44) was enriched for cell surface proteins *bam-2* and *sax-7,* as well as glutamate-gated chloride channel *avr-14,* all markers enriched in the HOA neuron ^51,52^.

Based on these gene expression signatures, we annotated each cluster accordingly. We predict that the PCA neurons correspond to the cluster labeled “Male 1,” based on high enrichment for *unc-103* ^53^ and *gar-3* ^54^ relative to the other male sensory neurons. However, the relatively low expression of ciliated neuron markers suggests that this cluster includes unciliated neurons, perhaps the related p.c.s. PCB and PCC neurons. The remaining predicted male-specific clusters that could not be confidently annotated were designated “Male 2” through “Male 12”. Together, we have identified 10/27 male specific neuron classes, with an additional 3 predicted classes, the p.c.s neurons. These data provide a substantially expanded transcriptional map of male-specific neural identities and a resource for the study of both unannotated and annotated neurons.

### Hierarchical clustering reveals functional organization and dimorphic relationships

Next, we asked how similar the neuron clusters are, based on gene expression profiles and using hierarchical clustering (Figure 2L). Consistent with prior observations showing that neurons cluster by type independently of developmental lineage^21^, our transcriptome-based diagram revealed that most neuron classes grouped predominantly by function (chemosensory, motor, etc.). Notably, known male-specific neurons such as CEM, HOB, and RnB clustered together, reflecting their shared molecular identity and roles in mating behavior. Similarly, RnA and HOA neurons also clustered together, but were distinct from their related ciliated HOB and RnB counterparts.

The dopaminergic ray type A neurons 5,7, and 9 clustered closely with the sex-shared dopaminergic CEP/ADE/PDE neurons. Importantly, most predicted male-specific or male-biased clusters also grouped together, reinforcing their sex-specific identity. Interestingly, the AWA, PHD, and SPV neurons clustered together, despite their anatomical and developmental differences. Given that SPV neurons are putative chemosensory neurons^55^ proposed to sense vulval pheromone and coordinate sperm release, and that PHD neurons were enriched for genes related to functions such as response to stimulus and localization, these features suggest that PHD neurons may have previously-unrecognized chemosensory functions.

### PHD sense female pheromone revealing a possibly novel sensory function

Building on the observation that PHD neurons clustered near known chemosensory neurons in our hierarchical analysis (Figure 2L), we investigated whether the PHDs might play a role in sensory behavior. Strikingly, the chemoreceptor for female volatile sex pheromone, *srd-1,* was not only highly expressed in AWA and ASI as expected^17^, but also in >40% nuclei within the PHD neurons (Table S2), suggesting the PHD might have a role in pheromone sensing. To test this hypothesis, we performed neuron-specific knockdown of *srt-18,* a gene highly enriched in PHD neurons, by using a pan-neuronal *sid-1* rescue strain in a *sid-1* mutant background (Methods). Adult males treated with *srt-18 RNAi* showed significantly impaired response to female pheromone (Figure 2M). Importantly, *srt-18* knockdown did not affect response to volatile odorants, such as benzaldehyde, nor did it impair locomotion in either sex (Figure S4C, D), suggesting the deficit is specific to male pheromone sensing. The PHD neurons were previously shown to mediate coordinated backward movement during mating^43^. Notably, a recent study confirmed *srd-1* expression in the PHD neurons, and revealed that expression of *srd-1* in both the AWA and PHD neurons serves to fine-tune volatile pheromone detection^56^. Our data further affirm the role of the PHD neurons in sensing more long-range mating cues, and suggest these sensory neurons may have a more complex sensory role, independent of *srd-1,* providing a previously-unrecognized sensory node in the male mating circuit.

### Sex-specific gene expression correlates with sex-specific behaviors

To identify male-enriched genes, we compared our male dataset to our previously published adult hermaphrodite neuronal snSeq dataset^21^. This analysis yielded 693 male-enriched genes, including 591 neuron-enriched targets detected in males and not detected in hermaphrodites (Table S4, Methods). Gene ontology analysis showed many of these genes were poorly characterized, with annotations limited to general cellular components. Among the better-characterized genes, there was significant enrichment for signal transduction-related functions, such as G protein-coupled receptor (GPCR) activity and phosphoprotein phosphatase activity (Figure 3A, Table S4). 7% of the male-enriched genes encoded C-type lectins. While about one-quarter of C-type lectins are present in the vas deferens^57^, we identified a distinct set of *clec* genes in CEM, HOB, and RnB neurons (Figure 3B, Table S5) where they may mediate male mating efficiency, similar to previously characterized lectins such as *clec-164*^7,33,57^.

**Figure 3:**
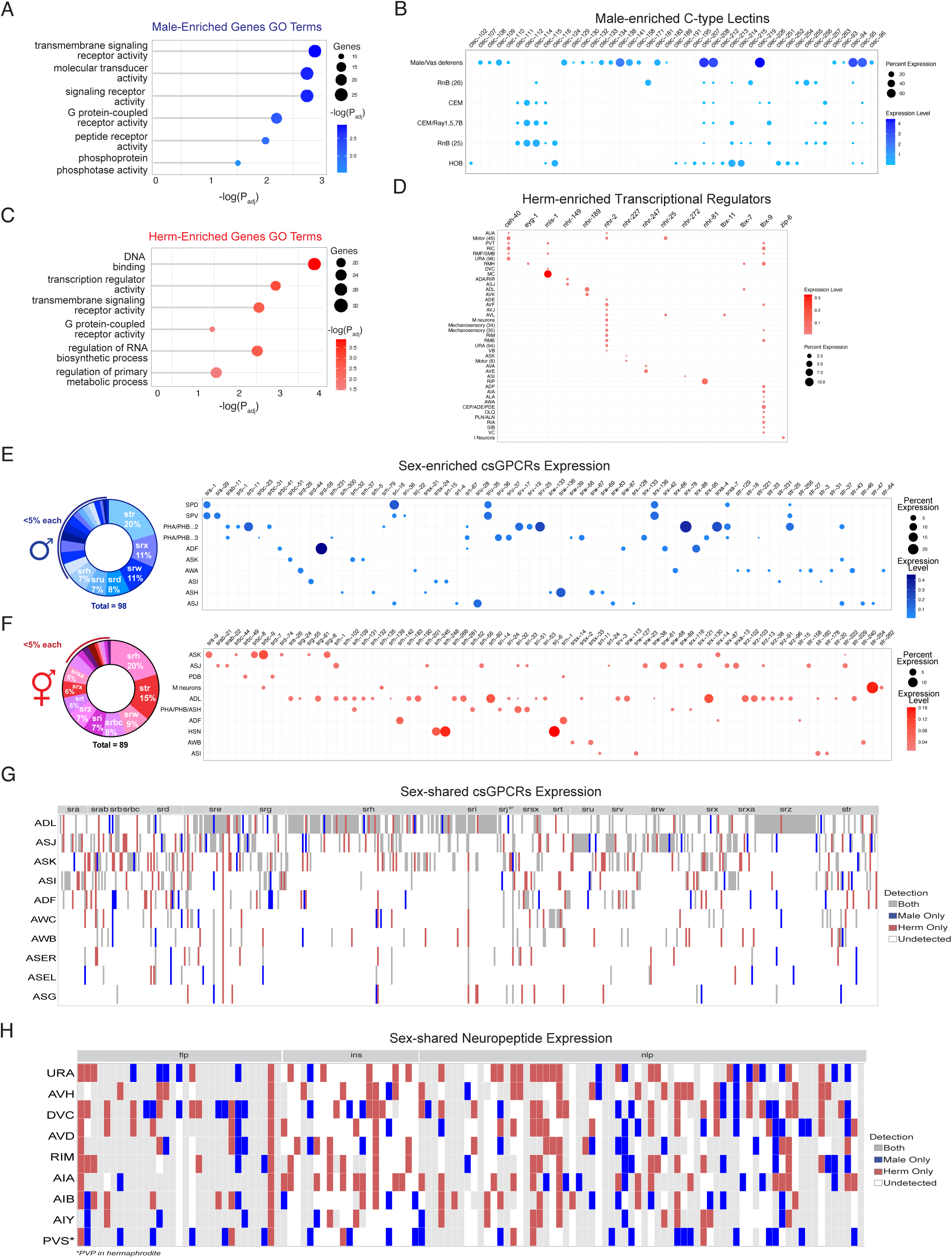
Sex-specific and shared targets reveal extensive transcriptional heterogeneity across neurons. **(A)** Gene Ontology (GO) terms associated with male-enriched genes not detected in hermaphrodites, generated using g:Profiler. **(B)** Expression of male-enriched C-type lectins in vas deferens and male-specific neurons. **(C)** Gene Ontology (GO) terms associated with hermaphrodite-enriched genes not detected in males, generated using g:Profiler. **(D)** Expression of herm-specific transcriptional regulators**. (E)** Expression of male-only csGPCRs and **(F)** herm-only csGPCRs, neurons with highest frequency of sex-specific expression shown. **(B,D,E,F)** Normalized average expression values and percentage of cells expressing the corresponding gene are shown. Numbers in parentheses correspond to cluster number associated with that neuron, used to distinguish clusters with the same name **(G)** Heatmap of sex-shared csGPCRs and **(H)** Neuropeptide expression between sexes. Genes expressed in the same cell type in both sexes are shown in gray. Those only in male are shown in blue, and only in hermaphrodites are shown in red, white space represents not detected. Genes present in >1% of cells with an average expression >0.001 in each neuron cluster are counted as expressed in that neuron.

Conversely, we identified 708 hermaphrodite-enriched genes, genes expressed in hermaphrodite neurons but absent in males (Table S4). These genes were similarly enriched for GPCRs but also showed significant enrichment for transcriptional regulators, including several nuclear hormone receptors (NHRs) and t-box transcription factors (TBX) (Figure 3C, D). Some targets such as *nhr-2* and *tbx-9* were broadly expressed, while others were limited to a few subsets of neurons (e.g., *tbx-11* and *zip-6*). *nhr* and *tbx* genes are known regulators of neuronal specification^58^ and sex determination during development^59^. However, the expression and regulation of these factors in adult neurons, and their roles in maintaining sex-specific neural states remains less well understood.

Given the critical role of GPCRs in sensory processing and the observed enrichment of sex-biased GPCRs, we next examined the dimorphic patterns of these chemoreceptors^60^. The *C. elegans* genome encodes roughly 1500 GPCRs, of which 1341 are putative chemosensory GPCRs^39^. These csGPCRs respond to pheromones, food-related cues, and repellents, driving behaviors like foraging and mate searching.

We identified 98 male-enriched csCGPRs (Table S6). Some, such as *srd-58* (ADF) and *srx-78* (PHA/PHB) were strong cell-specific markers. Notably, we confirmed the expression of known male-enriched GPCRs, including *sra-1*^39^ in SPV and SPD, *srr-7*^33^ in male CEM, HOB, and RnB neurons, and *srd-66* in the male CA7 neuron (Figure 2A-B, 3E, Table S2,S7). Some of these csGPCRs showed similar expression patterns in males as previously observed in larval hermaphrodites (e.g., *str-256* in AWA and *srh-37* in ASK^39^). Others, however, showed sex-enriched expression patterns. For instance, *srv-25* was expressed in PHA/PHB (Figure 3E) and male CP9 (Table S2), but was previously detected in L4 hermaphrodites^6^ and absent in adult hermaphrodites^21^. Similarly, *srh-300,* enriched in male ASH neurons, had been reported in larval-stage, but not adult^21^, hermaphrodite AVE^62^ and AVK^63^ neurons. *srw-87,* previously observed in embryonic ADF neurons^64^, was restricted to male ADF neurons in our dataset.

The expression patterns of these male-enriched GPCRs provided insights into neuronal specialization. Male-specific neurons such as SPV and SPD have several GPCRs in common, consistent with their homologous functional roles (Figure 3C, Table S6). Moreover, the majority of these male-enriched GPCRs, such as *srd-58*, are expressed in chemosensory neurons that are shared between the sexes, such as PHA/PHB, AWA, ADF, and ASK (Figure 3C). These neurons play established roles in mating behaviors. For example, PHA neurons, along with URY and PQR, regulate male reproductive drive through PDF-1 neuropeptide signaling^41^. The ADF^15^, and ASK neurons are key mediators of sex-dimorphic pheromone detection^65^. Therefore, the expression of specific csGPCRs in these neurons might suggest a role for these receptors in male-specific behaviors.

We also identified 89 csGPCRs that are expressed in adult hermaphrodites but not age-matched males (Table S4, 7). Here, we captured known markers for hermaphrodite-specific neurons, such as *srj-6,* in the HSN motor neurons (Figure 3F). Strikingly, the sex-shared ADL chemosensory neurons exhibited 32 hermaphrodite-enriched GPCRs compared to only two male-enriched ones (Table S4). Other chemosensory neurons, such as ASK and ASJ, also showed high enrichment. Many of these GPCRs had previously been identified in L4 hermaphrodite ADL neurons, such as *srx-130* and *srh-80*^6^, while others, such as *str-229* and *srz-38,* were newly reported in this study.

These data demonstrate that sexually dimorphic remodeling of chemoreceptor expression occurs in a developmental stage- and sex-specific manner, particularly in shared sensory neurons critical for reproductive behaviors.

### Sex-shared neurons display extensive molecular heterogeneity of shared transcriptome

To better understand how each of the sexes employs the same molecular machinery to regulate divergent phenotypic outcomes, we examined sex-enriched patterns of csGPCRs and neuropeptide expression in sex-shared neurons. First, we assessed the overlap of csGPCRs expression across both sexes. On average, ∼50% of the GPCRs displayed conserved expression patterns between the sexes within a given neuron (Figure 3G, S2E, Table S6). However, there were multiple examples of sex-enriched GPCR expression within a given neuron, or heterogeneity in expression of the same GPCR across different cell types between the sexes. For example, *sra-37* was detected in ASER and ADF in both sexes, but was hermaphrodite-enriched in AWC, ASI, and ASK, and male-enriched in ASEL. Chemosensory neurons ADL, ASJ, and ADF show highest expression of male- and hermaphrodite-biased GPCRs.

We next analyzed neuropeptide expression patterns, due to their function in modulating many sex-shared and specific behaviors in *C. elegans,* such as feeding, learning, and reproduction^41,66–71^. The *C. elegans* genome encodes over 100 neuropeptide genes, spanning the FMRFamide-related peptides (FLPs), neuropeptide-like proteins (NLPs), and insulin-like peptides (INS)^72–74^. We detected 133 neuropeptides shared between male and hermaphrodite^21^ neurons when comparing across the same threshold (Table S7). Several neuropeptides exhibit clear sex-biased expression, such as *flp-1*, *flp-8,* and *ins-30* (hermaphrodite-biased) and *nlp-43* and *flp-34* (male-biased). PVS (PVP in hermaphrodites), DVC, and URA neurons displayed the highest incidence of male-biased neuropeptides (Figure 3H). Chemosensory neurons including ASER, AWC, ASI, and ADL showed the highest prevalence of hermaphrodite-enriched neuropeptides (Table S7).

### Integrative analysis reveals extensive neuronal dimorphism and functional divergence

To further characterize sex-differential expression at the single-cell level, we integrated the male and genotypically-matched *him-8* hermaphrodite datasets, including 46,685 hermaphrodite neuronal nuclei (Figure S3). After processing, normalization, and unsupervised network clustering, we obtained 145 clusters, annotating 108 of 116 shared neurons classes (Figure 4A).

**Figure 4.**
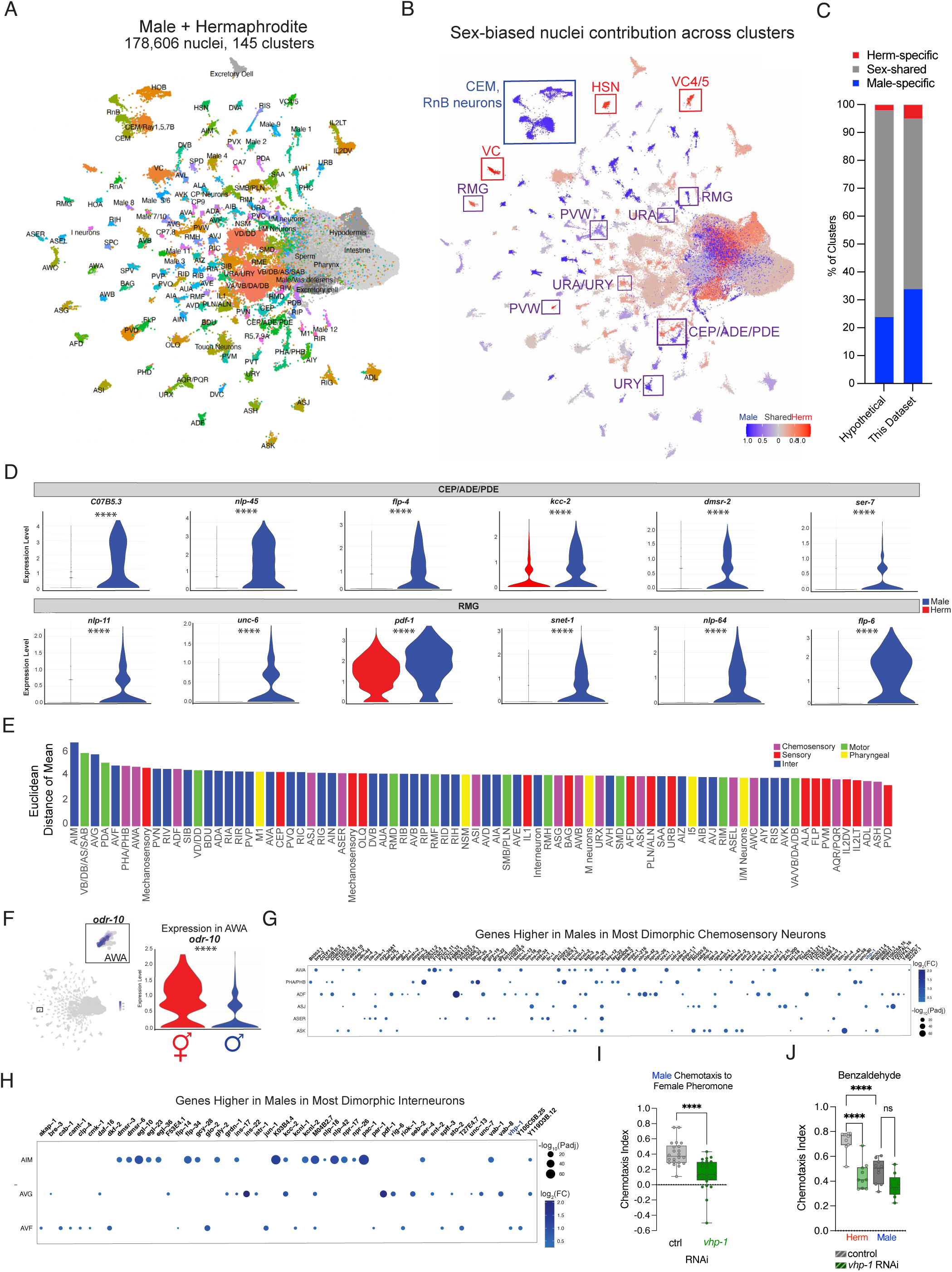
Sex-shared neurons reveal a highly dimorphic neuronal landscape. **(A)** UMAP of all 178,606 male and hermaphrodite nuclei that passed quality control across all biological replicates (4 male, 3 hermaphrodite), resulting in 145 distinct clusters. **(B)** UMAP projection of integrated male and hermaphrodite nuclei colored by sex bias. Each cluster is colored by normalized contribution of male vs hermaphrodite nuclei within each clustered (% male - % herm). Blue indicates male-biased, red indicates hermaphrodite-biased, and gray indicates balanced sex-shared groups. Sex-specific neurons (e.g., male CEM and RnB, hermaphrodite HSN and VC) are strongly enriched for the corresponding sex. RMG, URA/URY, PVW, and CEP/ADE/PDE are boxed in purple, highlighting sex-shared neurons segregating into distinct clusters. **(C)** Comparison of expected and observed distributions of sex-shared and -specific neuronal clusters. Stacked bar plots show the proportion of neuronal clusters that are male- specific/bias (blue), sex-shared (gray), and herm-specific/bias (red). The “hypothetical” bar represents expected distribution assuming complete recovery of all 294 sex-shared neurons, 93 male-specific neurons, and 8-hermaphrodite-specific neurons. The adjacent bar represents the distribution observed in our single-nucleus RNA seq dataset of both sexes. **(D)** Genes significantly higher in males in CEP/ADE/PDE and RMG neurons. Expression level density of male (blue) or hermaphrodite (red). Adjusted p-values from Wilcoxon Rank Sum test. **(E)** Euclidean distance of each neuron’s mean male vector and mean hermaphrodite vector. The top neuron subtypes with the largest distance are shown. **(F)** Feature plot of *odr-10* expression and violin plot of *odr-10* differential expression between sexes confirming upregulation in hermaphrodite AWA. **(G)** Genes higher in males (blue) in selected highly distant chemosensory neurons and **(H)** interneurons between the sexes. *vhp-1* is differentially expressed in ASK, ASJ, AVF (shown in blue)**. (I)** Knockdown of *vhp-1* in neuron RNAi-hypersensitive males impairs chemotaxis to *C. remanei* female pheromone. N= 3 biological replicates, 5-10 chemotaxis plates per replicate with ∼100-200 worms per plate. Unpaired, two-tailed Student’s t-test. **(J)** Knockdown of *vhp-1* attenuates benzaldehyde chemotaxis only in hermaphrodites. N = 2 biological replicates. One-way ANOVA with Bonferroni post hoc analysis. ****p<0.0001, ns: p > 0.05.

We next examined the normalized contribution of nuclei from each sex and biological replicate per cluster, generating a UMAP colored by this relative distribution to visualize sex-biased clustering (Figure 4B, Table S8). Sex-specific neurons, such as the CEM and RnB in males and the HSN and VC neurons in hermaphrodites, were >95% enriched for nuclei of the corresponding sex, validating the quality of our integration. Hierarchical clustering of the integrated dataset showed robust grouping of our predicted male-specific or male-biased neurons, including close clustering of Ray 5-,7,9A, HOA, Male 1, and RnA (Figure S5A), while the PHD neurons clustered with other chemosensory neurons. While most male-biased neurons were >90% enriched for male nuclei, 8 of 46 male-biased neurons showed 75-85% enrichment of male nuclei. Some of these were sex-shared neurons like the PHC, not identified in our previous hermaphrodite snSeq dataset^21^, likely reflecting technical differences in neuron recovery state. Based on these distributions, we classified clusters “sex-shared” (>30% nuclei from each sex), “hermaphrodite-specific” (<30% males), and “male-specific” (<30% hermaphrodite nuclei) (Figure 4C). This categorization resulted in 61% sex-shared, 5% hermaphrodite-specific, and 34% male-specific clusters, closely aligning with the expected distribution given known neuron numbers, 294 sex-shared, 8-hermaphrodite-specific, and 93 male-specific neurons (Figure 4C).

In fact, some sex-shared neurons were so dimorphic they clustered as separate cell types. For example, although the homologous gene expression and functional roles of CEP, ADE, and PDE - which comprise 8 sex-shared dopaminergic neurons with mechanosensory functions^31^ - cause them to cluster together, the sex-specific differences in expression cause the cluster to diverge. Similarly, the URA/URY neurons form a single cluster in hermaphrodites, but two distinct male clusters (Figure 4B, Table S2). Signaling-related genes, including neuropeptides (e.g., *nlp-45*, *flp-4, nlp-11, flp-6, nlp-64*), receptors (*dmsr-2, pdf-1, ser-7*), and signaling genes (*kcc-2, snet-1*), are some of the genes enriched in male neurons in these extensively dimorphic clusters (e.g., in CEP/ADE/PDE and RMG neurons; Figure 4D; Table S9). Another subset of genes was upregulated in hermaphrodites, including several neuropeptides, such as *flp-14, flp-5, nlp-46,* and *nlp-56* in RMG, and *nlp-10, nlp-6,* and *nlp-13* in PVW (Table S9).

To more precisely quantify the extent of molecular divergence between male and hermaphrodite neurons, we calculated the Euclidean distance (Table S10, Figure 4E). This analysis allows us to survey how different neurons are based on their expression profile. Here we only considered “sex-shared” clusters to ensure both sexes were sufficiently represented in each cluster, enabling biologically meaningful distance calculations. Clusters with fewer than 30% nuclei from one sex likely represent sex-specific (i.e., CEM) or highly dimorphic neuron types that cluster separately (i.e., URY), thus making direct distance comparisons less biologically meaningful. A few neurons such as AWA, enriched for 72% male nuclei, may be an exception to this rule and are possibly more readily isolated in one sex than another. Our data suggest that AIM interneurons were the most sexually dimorphic, in agreement with previous data demonstrating divergent gene expression and neurotransmitter identity between the sexes in this neuron class^75–77^. Other highly divergent neurons included AVG and AVF interneurons; VB/DB/SAB and PDA motor neurons; and AWA, PHA/PHB, and ADF, which were the most distant chemosensory neurons between males and hermaphrodites. Mechanosensory sensory neurons, including ALM and PLM, also showed high sex divergence.

We assessed the differentially-expressed genes between the sexes in these clusters, confirming expression of known sex-differential targets, such as upregulation of food chemoreceptor *odr-10* in hermaphrodites relative to males^16^ (Figure 4F, S3B). Similarly, we confirmed the upregulation of the canonical pheromone receptor^17^ *srd-1* in male AWA (Figure S5C). Overall, most differentially expressed genes across neurons were downregulated in males (Table S12). Thus, we posited that genes upregulated in males, like *srd-1,* may identify uncharacterized targets and their functional roles. Male upregulated genes include chemosensory and peptidergic GPCRs, neuropeptides, ion channels, and enzymes (Figure 4G, H, Table S12). There was also a global enrichment of ion channels, including *kcnl-1, kcnl-2, egl-23,* and *egl-26.* Over 100 genes were upregulated in the memory-associated AIM in males (Figure 4H, Table S12).

The MAPK phosphatase *vhp-1* showed higher expression in several male chemosensory and interneuron classes, including ASK, ASJ, and AVF (Figure 4G-H, Figure S5D, Table S12). Since ASK neurons play a role in pheromone sensing^26^ and are one of three sex-shared neurons required for sexual attraction in males, we hypothesized that *vhp-1* may play a role in pheromone sensing. Indeed, neuron-specific knockdown of *vhp-1* significantly impaired male attraction to female pheromone (Figure 4I). However*, vhp-1* is ubiquitously expressed across neurons, notably highest in chemosensory AWC neurons in which it is slightly more enriched in hermaphrodites (Figure S5E, Table S11). Therefore, we asked if *vhp-1* knockdown affected other functions in a sex-shared and/or specific manner. We tested the effect of *vhp-1* reduction on benzaldehyde chemotaxis in both sexes. Our data showed that *vhp-1* knockdown significantly reduces hermaphrodites’ ability to respond to benzaldehyde, but does not affect male chemotaxis to the same volatile odorant (Figure 4J). Yet knockdown of *vhp-1* did not affect pyrazine chemotaxis (AWA-regulated) or the animals’ movement in either sex (Figure S5F-G). Taken together, these results demonstrate that sex- and cell-specific regulation of sex-shared ubiquitously expressed genes can result in significantly divergent phenotypic outcomes.

## DISCUSSION

Here, we generated the first single-nucleus RNA-seq atlas of Day 1 adult *C. elegans* male neurons, alongside age- and genotype-matched hermaphrodite neuronal profiles. Using our recently-established snSeq protocol, we captured a comprehensive and high-purity dataset spanning both sex-shared and male-specific neurons. This atlas provides a powerful resource for identifying sex-specific and sex-shared programs underlying neuronal identity and behavior with single-cell resolution.

Through systematic annotation, we substantially expanded the molecular map of male-specific neurons, identifying markers for known male-specific classes and providing predictions for additional understudied neuron types. These markers will facilitate functional characterization of male neurons and annotation of both male and sex-shared neurons in future studies of sexual dimorphism in the nervous system.

Beyond male-specific annotation, our data revealed previously unrecognized relationships between sex-shared and sex-specific neurons. Our findings suggest that male-specific PHD neurons may have a role in pheromone sensing. Intriguingly, they also showed high transcriptional similarity with the sex-shared AWA chemosensory neurons, both enriched for genes related to chemosensory functions, including neuropeptides, csGPCRs, peptidergic GPCRs, and ion channels, such as TRPV channel *ocr-1.* This overlap suggests that sex-specific circuits partially re-purpose conserved molecular components. Intriguingly, OCR-1, which acts as a temperature sensor in AWA, is also upregulated in male AWA neurons compared to hermaphrodites, suggesting a similar role in PHD neurons, and possible sex-dimorphic thermosensory regulation in the sex-shared AWA. Human TRPV2, the OCR-1 ortholog, mediates mechanical nociception in the adult somatosensory system^78^ and its sex-differential regulation has been linked to higher prevalence of chronic pain in females. Thus, this overlap highlights the conservation of dimorphic sensory mechanisms across species.

We also uncovered extensive transcriptional heterogeneity across sex-shared neurons. Upon integration of our hermaphrodite and male single-cell data, we observed some sex-shared neurons, such as URY and dopaminergic neurons, diverged into distinct clusters, reflecting high transcriptional variability. While most sex-shared neurons clustered together as expected, we detected widespread sexually dimorphic expression patterns of GPCRs, neuropeptides, and ion channels. Among these were strong sex-and cell-type specific markers, including *srw-87,* specific to male ascarosides pheromone sensing ADF neurons, and *srg-8,* specific to hermaphrodite ASK neurons. Notably, *srw-87* was recently reported to be downregulated in male neurons with age^20^, raising the intriguing possibility that regulation of *srw-87* or other ADF gene targets may regulate the observed decline in pheromone sensing with age. While previous studies had reported isolated examples of sex-dimorphic neuropeptide^41^ and GPCR^39^ expression, these observation were often limited to single or few subsets of neurons. Our data now reveal that these differences are far more pervasive than previously appreciated, affecting diverse classes of neurons across the nervous system.

Our data reveal that ubiquitously expressed, sex-shared genes can exhibit striking sex- and cell-type specific functions. Although the MAPK phosphatase VHP-1 is broadly expressed across the nervous system, neuron-specific knockdown resulted in distinct, sex-specific behavioral phenotypes: impaired pheromone attraction in males and reduced benzaldehyde chemotaxis in hermaphrodites. These findings underscore the power of single-cell resolution to uncover functional divergence that would be masked in bulk analyses. While VHP-1 has been previously implicated in immunity and stress resistance^79,80^, its sex-specific roles had been unexplored. Interestingly, MAPK signaling has been shown to regulate dimorphic behavioral responses, such as higher pain sensitivity in female mice^81^ and increased response to alcohol induced liver injury in females with lower levels of MKP1^82^. Together, our findings, along with insights from mammalian systems, suggest a conserved role for MAPK signaling in regulation of neuronal functions in a sex-dependent manner, warranting further investigation into its contribution to sensory processing and stress resilience.

Taken together, our atlas not only expands the molecular annotation of male-specific neurons but also captures new sex-specific features and conserved differential expression patterns that may influence neurological function and behavior. By providing a comprehensive, high-resolution map of adult male and hermaphrodite neuronal transcriptomes, this resource will facilitate functional analyses of sex-specific and sex-shared circuits, help identify new candidate regulators, and advance the broader study of how sex influences nervous system function at the single-neuron level.

### Limitations of the Study

While we found that sequencing four male and three hermaphrodite biological samples was sufficient to resolve most of *C. elegans’* sex-shared adult neuron types (108 of 116), a few remained unresolvable. Additionally, we can only assess genes that we were able to measure in each cell type; we cannot comment on genes that may have been below the threshold of detection.

## Resource availability

### Lead contact

Further information and requests for resources and reagents should be directed to and will be fulfilled by the lead contact, Coleen T. Murphy.

### Materials availability

Worm strains generated in this study are available upon request. This study did not generate new unique reagents.

### Data and code availability

Single-nucleus RNA-seq data have been deposited at NCBI and are publicly available as of the date of publication. Accession number is PRJNA1195922 and is also listed in the key resources table. All other data are available in the main text or supplementary material.

This paper does not report original code.

Any additional information required to reanalyze the data reported in this paper is available from the lead contact upon request.

## Acknowledgments

We thank Christina DeCoste and Princeton FACS Core, Jennifer Miller, Jean Arly Vomar, Wei Wang, and the Princeton Genomics Core for their assistance, the *C. elegans* Genetics Center for strains, and members of the Murphy lab for suggestions on the manuscript.

## Funding

HHMI Gilliam Fellowship to KSM

The Simons Foundation (811235SPI) to CTM

China Scholarship Council (CSC) to YW

NSF GRFP to JSA (DGE-2039656)

NIH Office of the Director Pioneer Award to CTM (NIGMS DP1GM119167)

## Author contributions

Conceptualization: KSM, RK, JSA, YW, CTM

Methodology: KSM, RK, JSA, RK

Investigation: KSM, YW, JSA, RK

Visualization: KSM, JSA

Funding acquisition: CTM

’Project administration: CTM

Supervision: RK, CTM

Writing – original draft: KSM

Writing – review & editing: KSM, RK, CTM

## Declaration of interests

Authors declare that they have no competing interests.

## STAR Methods

### EXPERIMENTAL MODEL AND STUDY PARTICIPANT DETAILS

#### C. elegans growth and maintenance

All strains were maintained at 20C using standard methods for the duration of experiments^83^. *C. elegans* were maintained on plates made with high growth medium (HG: 3 g/L NaCl, 20 g/L Bacto-peptone, 30 g/L Bacto-agar in distilled water) with 4 mL/L cholesterol (5 mg/mL in ethanol), 1 mL/L 1M CaCl2, 1 mL/L 1M MgSO4, and 25 mL/L 1M potassium phosphate buffer (pH 6.0) added to the autoclave media after it has cooled to < 60C. Plates were seeded with a full lawn of OP50 *E. coli* (From the CGC) for *ad libitium* feeding. For RNA interference assays, animals were grown on HG seeded with OP50 until L4, then transferred onto HG made with 1 mM IPTG, 100 mg/L Carbenicilin, and 50 uM Floxuridine (FuDR) seeded with respective HT115 RNAi bacteria from the Ahringer Library^84^until day 2. Day 2 animals were then transferred to plates without FuDR seeded with the respective freshly inoculated HT115 RNAi bacteria. All behavior assays were performed on plates made with standard nematode growth medium (NGM: 3 g/L NaCl, 2.5 g/L Bacto-peptone, 17 g/L Bactoagar in distilled water) with 1 mL/L cholesterol (5 mg/mL in ethanol), 1 mL/L 1M CaCl2, 1 mL/L 1M MgSO4, and 25 mL/L 1M potassium phosphate buffer (pH 6.0) added to the autoclave media after it has cooled to < 60C. Hypochlorite synchronization was used to developmentally synchronize experimental worms, where gravid hermaphrodites were exposed to an alkaline-bleach solution (e.g. 7.5 mL sodium hypochlorite, 2.5 mL KOH, 41.5 mL distilled water) to collect eggs, followed by repeated washes with M9 buffer (6 g/L Na2HPO4, 3 g/L KH2PO4, 5 g/L NaCl and 1 mL/L 1M MgSO4 in distilled water)^83^.

#### C. elegans strains in this study

CQ760: *wqIs7 [Prgef-1::his-58::GFP] him-8(e1489) IV*

CQ830: *srd-66p::GFP;Pmyo3::mcherry;him-8(e1489)*

CQ828*: srg-1p::GFP;Pmyo3::mcherry;him-8(e1489)*

CQ859: *srh-217p::GFP;Pmyo3::mcherry;him-8(e1489)*

CQ860: *srt-18p::GFP;Pmyo3::mcherry;him-8(e1489)*

TU3595: *sid-1(pk3321) him-5(e1490) V; lin-15B(n744) X; uIs72 [pCFJ90(Pmyo-2::mCherry) + Punc-119::sid-1 + Pmec-18::mec-18::GFP]*

CQ826: *srg-14p::GFP;Pmyo3::mcherry;him-8(e1489)*

PB4641: *C. remanei*

### METHOD DETAILS

#### Filter design and efficiency testing

Adult males are shorter and more slender than hermaphrodites^22^, thus we capitalized on this size difference to separate the sexes. We removed existing meshes from 6-well tissue culture inserts (CellQart, item#: 9300002) and replaced them with 35 µm nylon meshes (BioDesign Inc. of New York, CellMicroSieves, product #: N35S) adhered to the inserts with a very thin film of epoxy resin (Loctite, epoxy adhesive, EA015) used along the insert border. Inserts are dried overnight, meshes are cut to shape, then sterilized with ethanol. The ethanol is allowed to evaporate, and the inserts are washed with M9 before filtering animals. We tested the efficiency of each individual filter using ∼200 µL of synchronized CQ760 hermaphrodites and males. We added 50 µL of animals to each individual filter and enough M9 to lightly submerge the animals. After visual inspection, each corresponding top and bottom fraction were then added to individual 15 mL conical tubes (8 total, 1 hermaphrodite- and 1 male-enriched tube/filter) and animals were allowed to settle. Then 5 µL of animals corresponding to each fraction was transferred to NGM plates seeded with 150 µL of OP50, animals were allowed to spread then paralyzed with 7.5 % sodium azide to facilitate counting. The number of males and hermaphrodites on half of each plate were counted. We then calculated the percent filtering efficiency. It is important to note that filters may tear, decreasing their efficiency, but this is easily visualized upon inspection of 6-well plates under a standard microscope prior to downstream processing. Filter efficiency is calculated as follows:

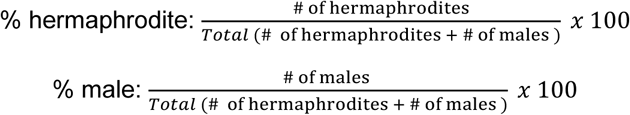

#### Neuronal nuclei isolation

*C. elegans* neuronal nuclei were isolated as previously described^21^ with minor modifications. About 0.8-1 mL of Day 1 synchronized hermaphrodites and males (CQ760) were washed from HG plates, then washed 3X times with M9 to get rid of excess OP50. Animals were then transferred to 6-well inserts with 35 µm nylon filters submerged in M9 to separate both sexes. We use 4 filters for each replicate. Roughly 150-200 µL of animals are added to each filter and allowed to filter for ∼5 min. Upon inspection of the top fraction (hermaphrodite-enriched) and bottom fraction (male-enriched), the respective fractions are pipetted into fresh 15 mL conical tubes for downstream nuclei isolation.

First, hermaphrodites were Dounce homogenized in lysis buffer 30-50x and males 50-90x to break open the cuticle. Briefly, after chemo-mechanical lysis, the pellet was resuspended in 500 µL wash buffer, Hoechst stained (1:10,000 dilution; Molecular Probes Hoechst 33342, Thermo Fisher, Cat. # H3570) was added to each sample, and samples were passed through 5 μm syringe filters, directly into FACS tubes. Samples were incubated for at least 5 min on ice prior to FACS.

Nuclei positive for both Hoechst and GFP+ were sorted using a 70 μm nozzle and a flow rate of 3 on a BD Biosciences FACSAria Fusion sorter into a 1.5 mL low bind microcentrifuge tube containing collection buffer (500 μL of 0.5% BSA + 1.5. U/μL RNase Inhibitor). The instrument was washed with bleach between samples. We performed 3 biological replicates for hermaphrodites and 4 biological replicates for males to increase the number of male nuclei. For the 4^th^ male biological replicate samples were split into two technical replicates by splitting ∼300 µL of filtered males into two aliquots and following all described downstream nuclei isolation steps.

#### Library preparation, sequencing, alignment, and QC

After FACS, samples were centrifuged gently at 1000 x g for 5 min at 4°C. The gentle speed, while important to keep the nuclei intact, leaves many of them behind in the supernatant. The supernatant was removed, and nuclei were resuspended in 20 μL of collection buffer. Single nuclei suspension samples were then provided to the Princeton Genomics Core for 10X Genomics Chromium X barcoding, cDNA amplification, library preparation and Illumina next generation sequencing, as described in St. Ange and Weng et al. 2024^21^. After Illumina sequencing, reads were aligned using CellRanger version 7.1.0. SoupX was used to remove ambient RNA contamination on Cell Ranger output files, and calculated contamination fractions were between 0.12 - 0.50 for all samples.

Next, we assessed quality control metrics to give us insight into the depth, complexity, and overall quality of our data. Average genes/cell, average UMIs/cell, and number of cells per cluster were all assessed. Violin Plots of genes per cell were generated to determine cutoffs: the lower bound was 100-300 features/cell for each sample (to remove damaged nuclei and empty droplets) and the higher bound was between 750 and 1500 features/nucleus (to remove doublets) depending on the sample. Data outside of these cutoffs were excluded from further analysis.

#### Normalization, Integration, and Clustering

We used the Seurat package single cell genomics pipeline for normalization, integration and clustering. We first merged all of the replicates. Next, we normalized the data using single cell transform (SCT), which normalizes single cell data by fitting genes to a negative binomial distribution. We generated an elbow plot of the principal components (PCs) to determine the number of PCs to include in dimensional reduction, and we used the Louvain algorithm for unsupervised network clustering. We tested different resolutions: 0.6 - 2.4 in 0.2 intervals. Ultimately, we clustered the data using 150 PCs at a clustering resolution of 1.4. This resulted in 110 clusters for the male data processed on its own, and 145 clusters for the male and hermaphrodite data processed together.

#### General Cluster Labeling/Cell Type Analysis

Cluster annotation for the male-specific transcriptome was performed using a combination of systemic and manual approaches as described in St. Ange & Weng et al. 2024^21^, with some additional curation used for understudied male-specific clusters. We first focused on only the male biological replicates to characterize the male transcriptome.

First, we used the ‘FindAllMarkers’ function in Seurat to identify cluster-specific markers. These markers corresponded to genes showing significant differential expression (log_2_FC > 0.25) within each cluster compared to the average expression across all clusters and were expressed in >25% of cells in the corresponding cluster.

We then compared the identified cell markers to a curated list of known markers for each neuron type classification (Table S1). This list comes from existing sequencing data and literature. Two statistical tests were used for this comparison: 1) A hypergeometric test, assessing the Bonferroni-corrected p-value of the overlapping genes (genes present in both the cell markers list and the curated list). Clusters demonstrating a significant overlap (p < 0.01) with known anatomy markers were considered for the corresponding neuron type. 2) An Area-Under-the-Curve (AUCell) algorithm^85^, which accounts for level of expression of a given gene set within the respective cell. Gene set lists were generated based on the known markers associated with a given neuron type and ranked by expression level in the cell in question. Using these rankings, we can calculate the AUC value of each gene set in each cell. To assign neuron types to cells, we generated a histogram of AUCell values and applied a threshold, manually adjusting the threshold in some cases to ensure approximately 5% of cells were assigned to each neuron term. Subsequently, we assembled these cells into clusters and determined the percent assigned to each neuron type. Neuron types with the highest 10 percentages were assessed for annotation. The agreement of these two statistical tests allows us to annotate clusters with neuronal identities in an unbiased way.

To add confidence to our annotations, we performed the hypergeometric test on not only the curated list, but also using the neuron type markers reported by St. Ange & Weng et al. 2024^21^. To annotate a given cluster, we used the results from the two hypergeometric tests, and the AUCell algorithm. If all three methods aligned, we regarded this as the final annotation. In case of disagreement, we examined gold-standard markers associated with that neuron and made a manual decision.

#### Annotation of Male Specific Neurons

For most male-enriched neurons (>70% nuclei stemming from a male sample when clustered with hermaphrodite biological replicates), markers are sparse, thus the statistical tests rarely reflect neuronal identity. First, we chose markers in our data that appeared cluster specific to generate fluorescent reporters to inform neuronal annotation. In instances where a male-enriched neuron could not be annotated through statistics or fluorescent reporters, it was labeled as closely to its neuron class as we could: male inter/motor “x”/male sensory. The results of both mechanisms and the final annotation are summarized in supplemental Table S1.

#### Annotation of Hermaphrodite and Male Clusters

To annotate our hermaphrodite and male integrated dataset, we used the markers identified in our male-only analysis to repeat the hypergeometric test. We also repeated the hypergeometric test original list of known markers and the markers reported by St. Ange & Weng et al., as we well as the AUCell test for further confirmation. For sex-shared clusters where all tests aligned, we regarded this as the final annotation. For male-enriched clusters (>70% nuclei stemming from a male sample) where tests didn’t agree, we deferred to the annotation based on the markers from the male-only data analysis. For hermaphrodite-enriched clusters, we deferred to our standard approach and manual curation of gold-standard markers. The results of all tests and the final annotation are summarized in supplemental (Table S8).

#### Hierarchical clustering

We used the normalized expression level of each gene in a given cluster to obtain average gene expression vectors, then calculated the Euclidean distance matrix between clusters. Lastly, we hierarchically clustered using the “hclust” function with the “complete” linkage method.

#### Threshold Setting for Expressed Genes

Similarly to CeNGEN^7^ and St. Ange & Weng et al. 2024^6,21^, we applied expression-level thresholds to the genes in our male-only dataset before assessing enrichment of a given gene in a particular cell type. We applied 4 thresholds that require a varying percent of expression of a gene within a cluster (0.5, 1, 1.5, or 3) and a minimum average normalized expression value (0.001) (Table S3). The same thresholds were applied to the hermaphrodite and male integrated dataset (Table S11).

#### Differential Gene Expression Analysis

Differential expression analysis was conducted as previously described^21^. Briefly, we used the FindMarkers function in the Seurat package and performed the Wilcoxon Rank Sum Test method on each neuron type, comparing male and hermaphrodite cells within the same cluster. Genes with a minimum percentage of expression in 10% of the cells were analyzed, and the significantly differentially expressed genes were identified if their log_2_(FC) > 0.25 or < −0.25, and adjusted p-value < 0.05.

Sexually dimorphic neurons that separated during unsupervised network clustering were merged into a singular Seurat object before differential expression. These neurons are the CEP/ADE/PDE, RMG, and PVW. We used the same statistical cutoffs for these neurons.

Heatmaps and dotplots were generated with the ggplot2 package in R.

#### Comparison to Hermaphrodite Data

For comparison with the St. Ange and Weng et al dataset, we asked what genes were detected in our male-only dataset threshold 2 that were not detected in their hermaphrodite threshold 1 (least stringent). St. Ange and Weng et al use the same thresholds as those in this study. In this manner we were able to identify male-enriched genes. The same approach was used to identify hermaphrodite-enriched genes. To analyze sex-shared candidates, we compared the N2 WT threshold 2 with our male threshold 2. Since we cannot compare expression levels across these different genotypes, we simply asked what genes were detected in each cluster for each sex based on the 1% threshold. Only sex-shared neurons were analyzed in this case.

#### Gene Ontology Analysis

Upon comparing the male threshold 2 genes to the list of wild-type Day 1 hermaphrodite threshold 1 genes, we identified a list of 693 male-enriched genes and 708 hermaphrodite-enriched genes (Table S4). We then used g:Profiler ^86^ to perform functional enrichment analysis on this set of genes and chose significantly enriched molecular functions, biological processes, and/or cellular components (Bonferroni adjusted p-value < 0.05). Similarly, we identified PHD- enriched genes using the FindMarkers function in Seurat and performed functional enrichment using g:Profiler.

#### GFP reporter construction

Promoters upstream of the ATG start site of target genes were PCR amplified, and restriction enzyme cloned into the pPD95.75 Fire vector (Addgene). The promoter region for each construct was 5 kb upstream of the start site or up to the stop codon of the nearest coding gene, whichever criteria was met first. Promoter::GFP plasmids were injected into N2 animals at 25 ng/µL, with 1 ng/µL of *Pmyo3::mCherry* as a co-injection marker.

#### Fluorescent microscopy

Fluorescence microscopy was performed on the Nikon AXR confocal microscope using 60X magnification. 20X magnification was used to visualize pan-neuronal expression. Animals were synchronized and grown on HG plates until day 1. Day 1 animals were paralyzed with sodium azide on agar pads. We used 488 excitation to image our GFP reporters. Images were collected using a Z-stack with 0.4 uM increments. Images are processed with FIJI. Processing includes maximum intensity projection, brightness/contrasts adjusting, de-speckling, rotation, thresholding, splitting, and merging color channels. We evaluated expression of each construct in both sexes in at least 5 animals. Images were taken in regions where expression was observed, and 2-3 images were taken of hermaphrodites lacking expression of male-specific target genes. DiI staining was performed by adding 5uL of 2 mg/mL DiI into 1 mL of M9 and allowing animals to gently mix in this solution for 2 hrs prior to washing with M9, then imaging. Here we image using 488 and 561 excitation for GFP and mCherry. Hermaphrodites and males imaged for size measurements were synchronized, paralyzed with sodium azide on agar pads, imaged using a 60X objective, then measured using the FIJI measurement tool.

#### Chemotaxis assays

Male pheromone preference assays were performed as previously described^20,26,87^ using *sid-1* mutant males with a pan-neuronal *sid-1* rescue for neuronal-specific knockdown upon RNAi treatment (described above). We used *C. remanei* true female pheromone since it has been shown to elicit a more robust response than *C. elegans* hermaphrodite pheromone^17,87^. *C. remanei* L4 females were picked onto freshly seeded NGM plates to ensure that they were unmated. Once females reach day 4, they were picked into M9 buffer with a concentration of 10 worms/100 µL M9. Animals are kept in M9 overnight to obtain a pheromone rich supernatant. The supernatant is then isolated for experiments. Pheromone may be kept at −20 °C for up to a month using single freeze-thaw cycles but works best fresh. For chemotaxis assays, 1 µL 7.5% sodium azide was added to each spot to paralyze the worms at each spot. Then, 2 µL of pheromone and M9 buffer control was spotted 5 cm apart on an unseeded 60 mm NGM plate. Males were washed 3x and 5 µL of lowly-dense pellet of males was added in-between the two spots. Then, males were given 1 hr to chemotaxis to either spot. Chemotaxis toward 1% benzaldehyde (Millipore Sigma Cat. #B1334-100G) in ethanol or 10 mg/mL pyrazine (Sigma-Aldrich Cat. #P56003-5G) in ethanol was accessed using standard conditions and performed on 10 cm NGM plates ^88^. Chemotaxis index is calculated as follows:

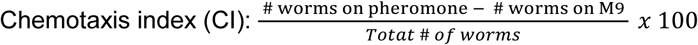

#### Euclidean distance analysis

First the data was subset into each cluster, and further subset by sex (i.e. males and hermaphrodites) using the Seurat package. Only clusters categorized as sex-shared were analyzed, as marked by roughly ∼30% or more nuclei from each sex present in that cluster. For males and hermaphrodites in each cluster, we generated a representative PC vector by averaging the column means of PCA embeddings from all cells in that cluster and genotype.

Then, we calculated the Euclidean distance between the PC vector of male and hermaphrodite in each cluster. We ranked the distance from the largest to the smallest.

### QUANTIFICATION AND STASTISTICAL ANALYSIS

Unpaired, two-tailed Student’s t-test was performed to compare chemotaxis with only two conditions. Wilcoxon Rank Sum test’s adjusted p-values are used for the identification of differentially expressed genes. Experiments were repeated on separate days, using separate independent populations, to confirm that results were reproducible. Prism 9 software was used for all statistical analyses. Software and statistical details used for RNA sequencing analyses are described in the method details section of the STAR Methods. Additional statistical details of experiments, including sample size (with n representing the number of chemotaxis assays performed for behavior, RNA collections for RNA-seq, and the number of worms for microscopy), can be found in the figure legends.

### Supplementary Table Legends

**Table S1. Male cluster annotation results related to Figure 1**

A curated list of previous known markers for each neuron type is listed in the “Neuron Type Markers” tab. A list of markers based on the St. Ange and Weng et al, 2024 Day 1 adult hermaphrodite data is listed as “Neuron Markers based on N2 Day 1”. Both were used to inform annotations in this work. Enriched genes for each identified cluster are shown next (pct1 = cluster of interest, pct2 = all clusters). We defined neuronal identity of each cluster based on hypergeometric test and AUCell test results, incorporating both list of markers provided. The test results and final annotation of each cluster are shown in the “Final Male Cluster Labels” tab. We also provide a list of male-specific neurons describing the neuron-type and whether they were identified in this study.

**Table S2. All genes and thresholds in male data, related to Figures 1-3**

Average normalized expression data for all genes detected in our male dataset separated into 4 thresholds (0.5%, 1%, 1.5%, and 3% cutoffs) outlined in Methods. There is a summary tab detailing the spread of average normalized expression values throughout each threshold.

**Table S3. Ciliated marker genes assessed in predicted male-specific neurons**

Expression of 10 ciliated sensory neuron marker genes and their average normalized expression in each male-specific cluster, as well as the percent of nuclei expressing the corresponding marker.

**Table S4. Male- and hermaphrodite-enriched genes related to Figure 3**

Average normalized expression values and percent expression for the 693 male-enriched genes and 708 hermaphrodite enriched genes referenced in Figure 3. GO terms generated from each list using g:Profiler are also provided.

**Table S5. Male-enriched C-type lectins related to Figure 3B**

Average normalized expression values (SCT) and percent expression (PCT) for the 47 male-enriched C-type lectins referenced in Figure 3B.

**Table S6. Male- and hermaphrodite-enriched and shared GPCRs, related to Figure 3E-G**

Average normalized expression values (SCT) and percent expression (PCT) for the 98 male-enriched GPCRs and 89 hermaphrodite-enriched GPCRs referenced in Figure 3, as well as expression of 596 sex-shared GPCRs assessed. Note that expression data is only shown for the sensory neurons referenced in Figure 3. However, all male expression data can be found in Table S2 and hermaphrodite data in St. Ange and Weng et al., 2024 Table S4.

**Table S7. Analysis of sex-shared neuropeptides related to Figure 3**

Overview of sex-shared and sex-enriched neuropeptides referenced in Figure 3H. Average normalized expression for each gene is shown separately for each sex.

**Table S8. Male & hermaphrodite integrated cluster annotation results related to Figure 4**

A list of markers generated based on enriched genes in each annotated cluster in our male-only analysis (shown in Table S1). Enriched genes for each identified cluster are shown (pct1 = cluster of interest, pct2 = all clusters). We defined neuronal identity of each cluster based on hypergeometric test and AUCell test results, as well as annotation strategies outlined in the text. The test results and final annotation of each cluster are shown in the “Male & Herm Cluster Annotation” tab.

**Table S9. Differentially expressed genes in sex-biased clusters shared between the sexes**

Male vs. hermaphrodite differential expression results from Wilcoxon Rank-Sum test of sex-biased neurons clustering separately between the sexes. Clusters were merged into a single Seurat object for analysis (Methods). Significantly differentially expressed genes within each cluster are shown in individual tabs with their gene name, p-value, average log2Fold-change (male/hermaphrodite), pc1 (male), pc2 (hermaphrodite), and adjusted p-value.

**Table S10. Euclidean distance measurements related to Figure 4E**

Euclidean distance between the PC vector of male and hermaphrodite in each cluster. Only sex-shared clusters were analyzed.

**Table S11. All genes ad thresholds in male and hermaphrodite integrated data related to Figure 4**

Average normalized expression data for all genes detected in our male and hermaphrodite integrated dataset separated into 4 thresholds outlined in Methods and split by sex. The summary tab details the spread of average normalized expression values throughout each threshold and contains a color-coded list of “sex-shared” and “sex-specific” neurons for reference along with the corresponding cluster numbers for each annotated cluster.

**Table S12. Male vs. hermaphrodite differentially expressed genes related to Figure 4**

Male vs. hermaphrodite differential expression results from Wilcoxon Rank-Sum test of every cluster. Significantly differentially expressed genes of each cluster are shown as a separate tab for each annotated cluster, with their gene name, p-value, average log2Fold-change (male/hermaphrodite), expression percentage pc1 (male) and pc2 (hermaphrodite), and adjusted p-value. Order of tabs corresponds to cluster numbers referenced in Table S8 and S13. Notably, differential expression values in sex-specific clusters are not informative, as these clusters are predominantly composed of neurons from a single sex. This prevents meaningful comparisons due to artificially inflated or absent differential signals. Instead, it is advised to refer to the expression data in Table S11 for gene expression information in sex-specific neurons.

## Key resources table

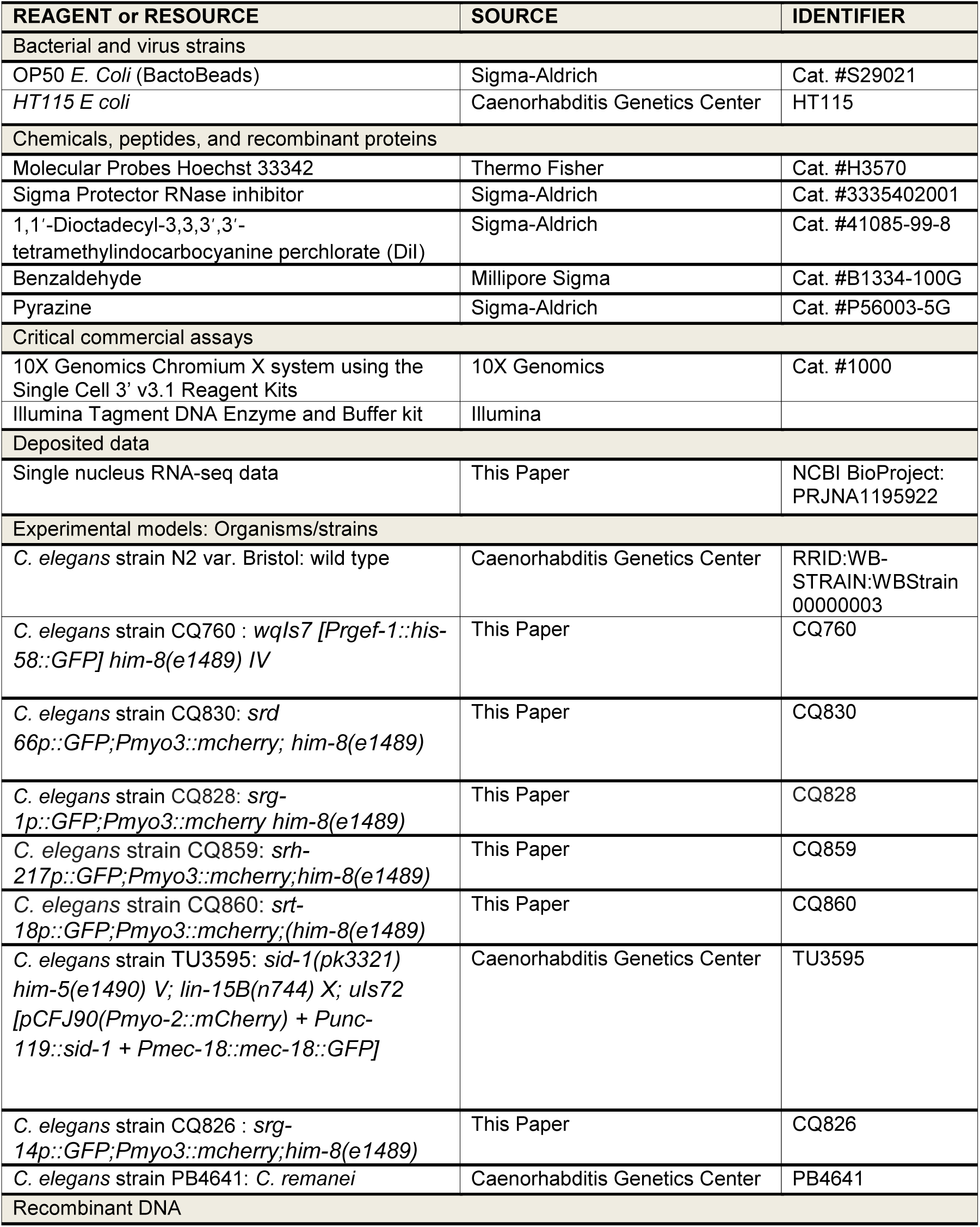
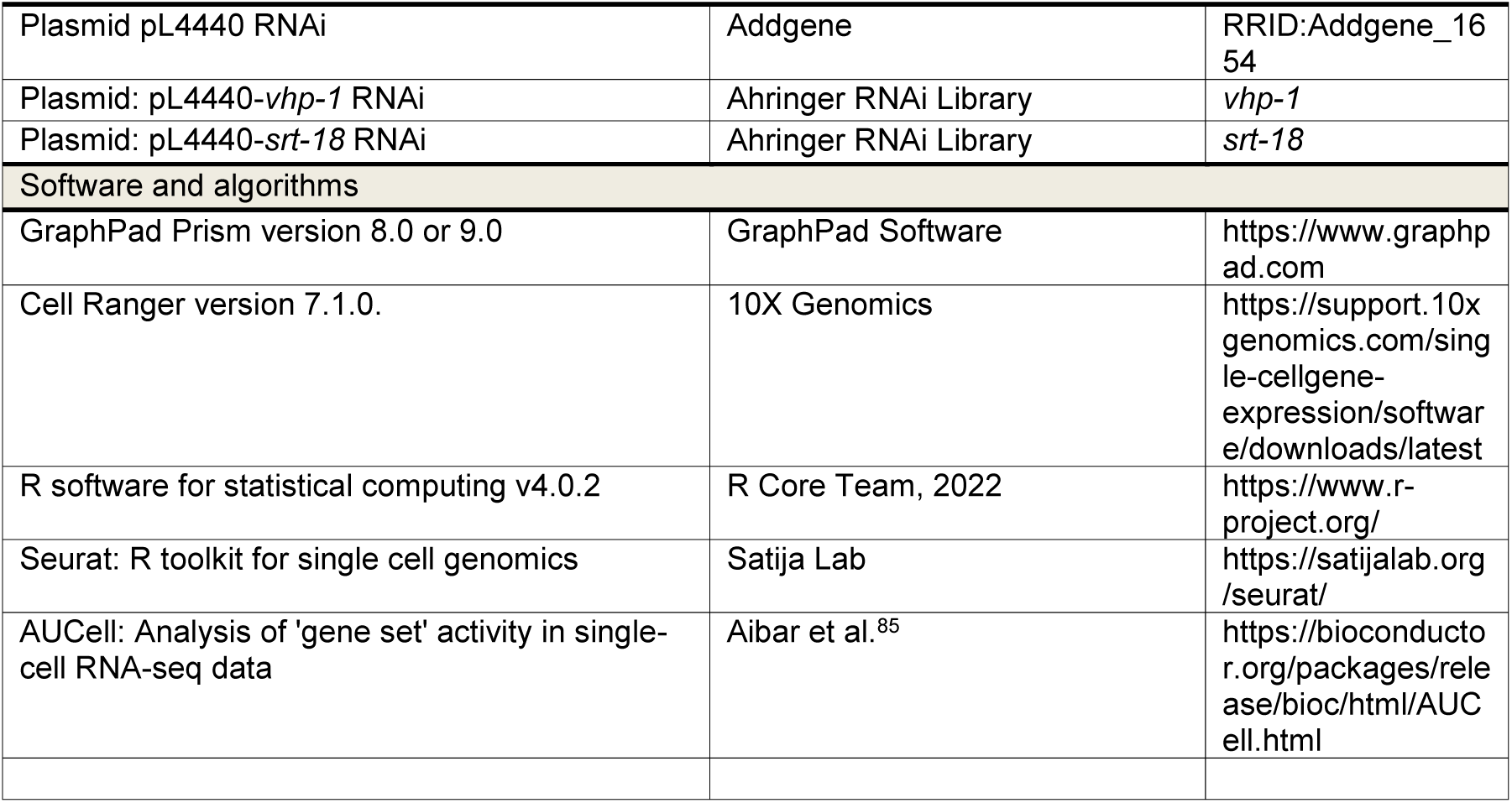

---

**Supplementary Figure 1:**
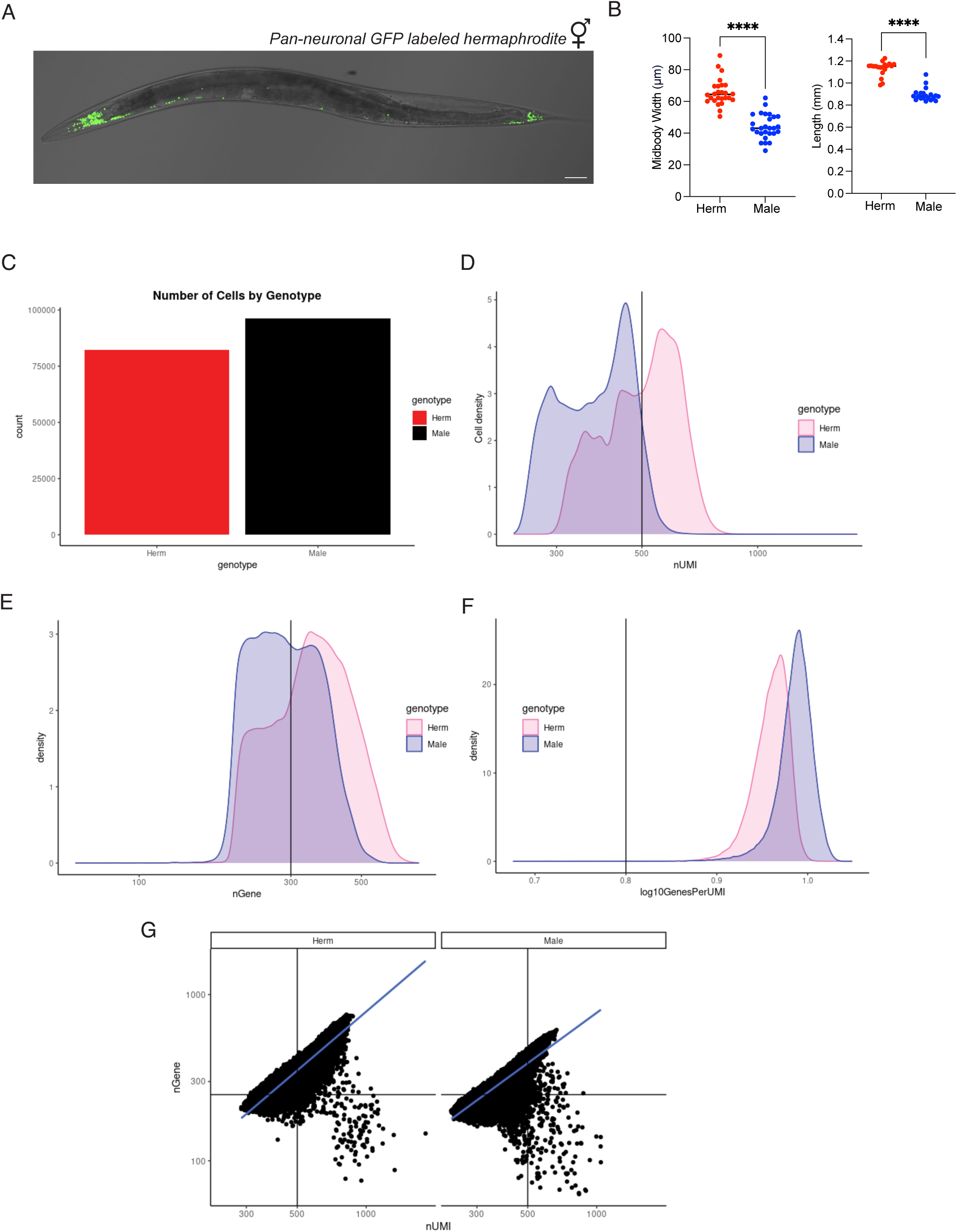
Quality control metrics for single nucleus RNA-seq. **(A)** Representative image of pan-neuronal GFP tagged *him-8* hermaphrodites (*Prgef-1::his-58::GFP;him-8*). Shows clear labeling of nuclei and less neurons in the tail compared to male. Scale bar: 50 µm **(B)** Midbody width(µm) and length(mm) comparisons between Day 1 males (blue) and Day 1 hermaphrodites (red). Worm widths were measured at the midbody using FIJI. Each dot represents a single animal. Hermaphrodites have an average width of 60 µm while males are 40 µm wide on average, allowing them to squeeze through 35 µm filters. Unpaired two-tailed Student’s t-test, ****p < 0.0001. **(C)** Number of cells across all replicates for each sex. **(D)** Number of UMIs per sex. **(E)** Number of genes per cell density. **(F)** Genes per UMI relative to cell density. **(G)** Number of genes relative to the number of UMIs across both sexes.

**Supplementary Figure 2:**
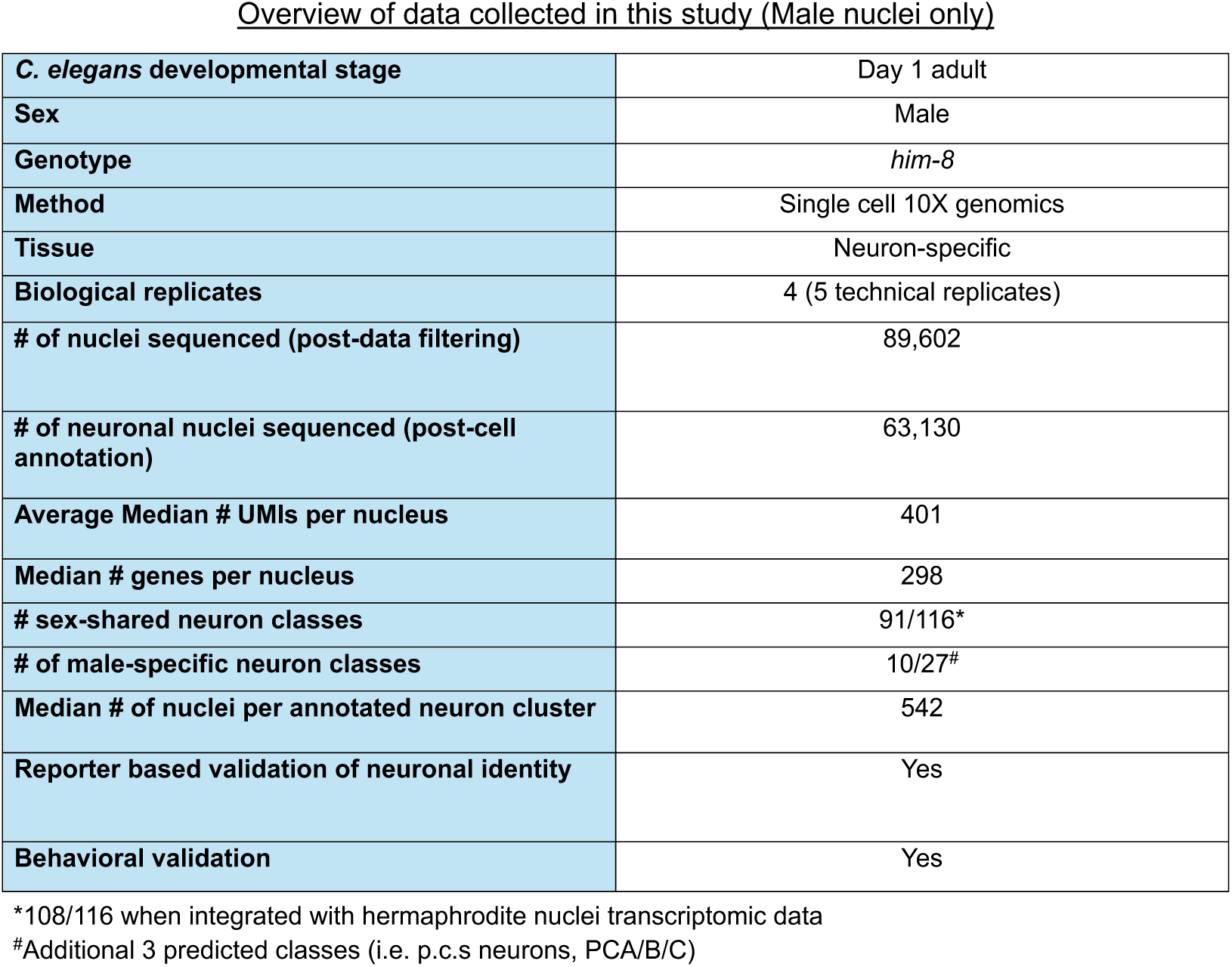
Overview of male snSeq data. Provides summary information for the 4 biological male replicate samples used for analysis of the male neuronal transcriptome. This includes developmental stage, genotype of the animals used, sequencing method, number of nuclei sequenced (post-data filtering and post-cell annotation), and median UMI/genes per nucleus.

**Supplementary Figure 3:**
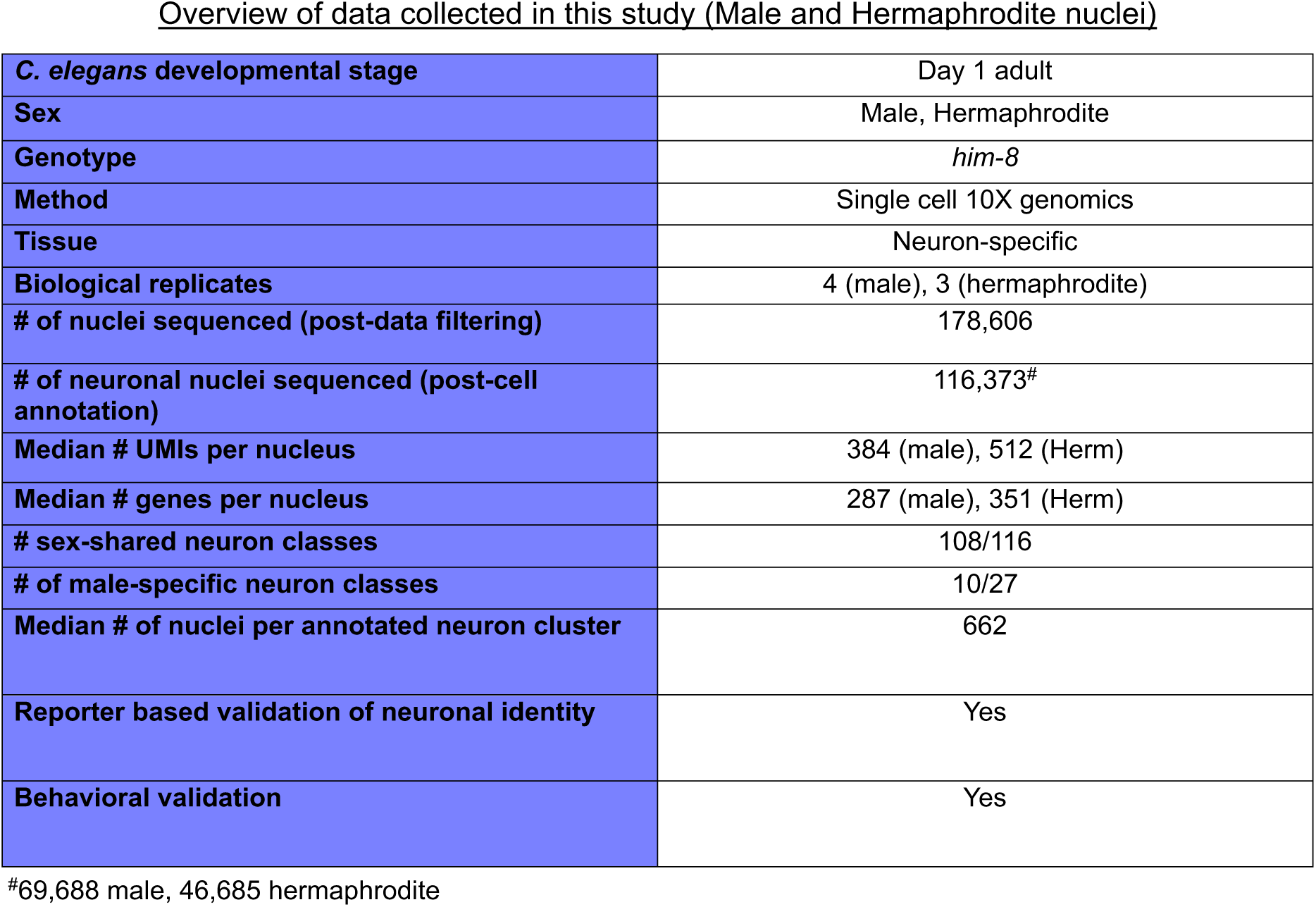
Overview of male and hermaphrodite snSeq integrated data. Summary information for 4 biological male replicates and 3 biological hermaphrodite replicates sequenced for analysis of the male and hermaphrodite neuronal transcriptomes. This includes developmental stage, genotype of the animals used, sequencing method, number of nuclei sequenced (post-data filtering and post-cell annotation), and median UMI/genes per nucleus.

**Supplementary Figure 4:**
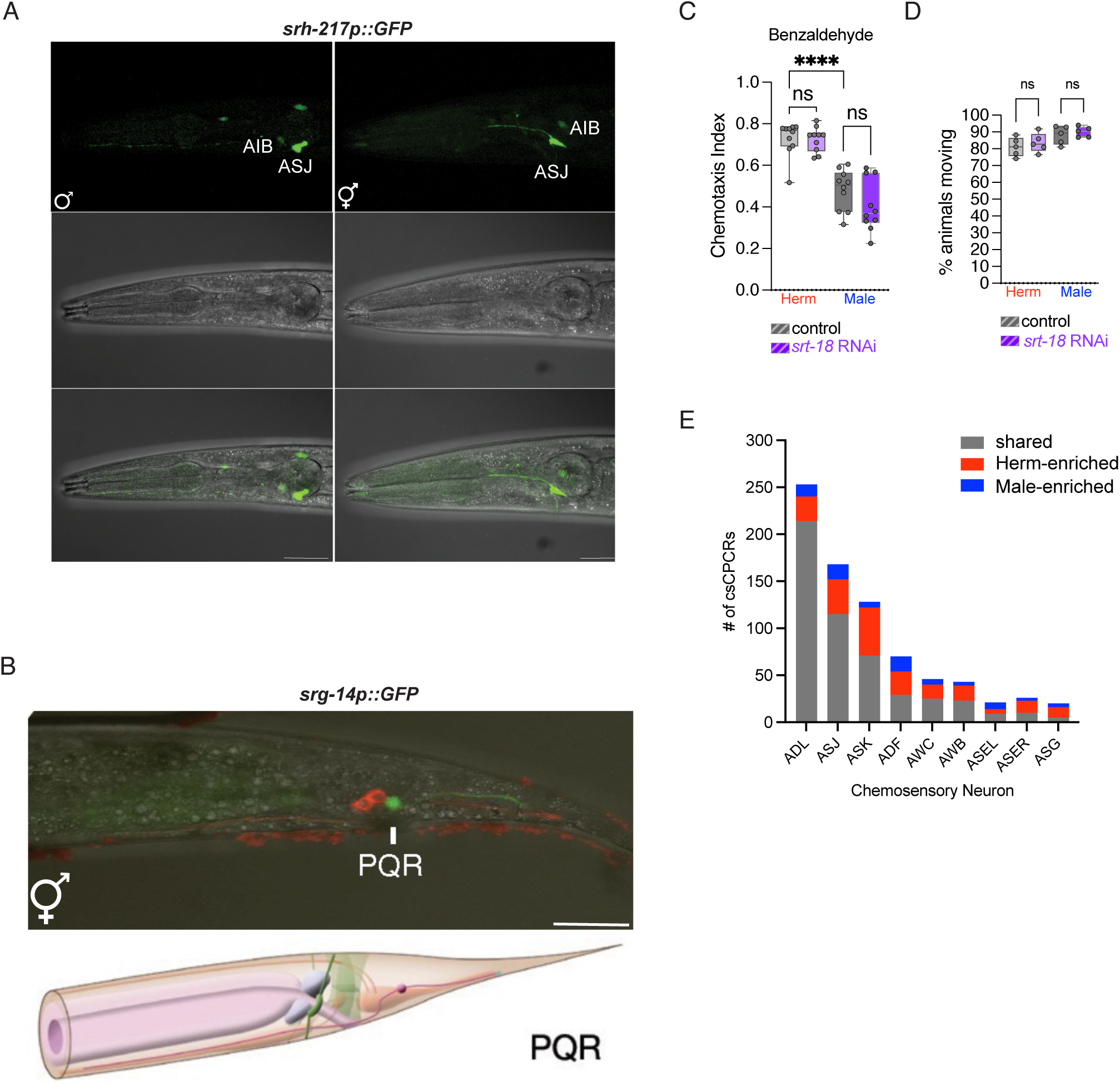
Reporter expression patterns, behavioral effects of *srt-18* knockdown, and chemosensory GPCR distribution. **(A)** Imaging of *srh-217p::GFP* shows expression in ASJ and AIB in both males and hermaphrodites. **(B)** Merge image of brightfield, DsRed and GFP channel after DiI staining of Day 1 hermaphrodites. DiI stains phasmid PHA and PHB and serves as reference point validating nearby location of PQR. (A, B) Scale bar: 25 µm. **(C)** Benzaldehyde chemotaxis is unaffected by *srt-18* neuron-specific knockdown. Males show reduced preference to benzaldehyde (1%) compared to hermaphrodites as previously observed. (D) *srt-18* RNAi knockdown does not affect animals’ ability to move compared to vector control. One-way ANOVA with Bonferroni post hoc analysis ****p<0.0001, ns: p > 0.05. **(E)** Stacked bar plot showing distribution of sex-shared and sex-biased GPCRs in chemosensory neurons presented in figure 3G.

**Supplementary Figure 5:**
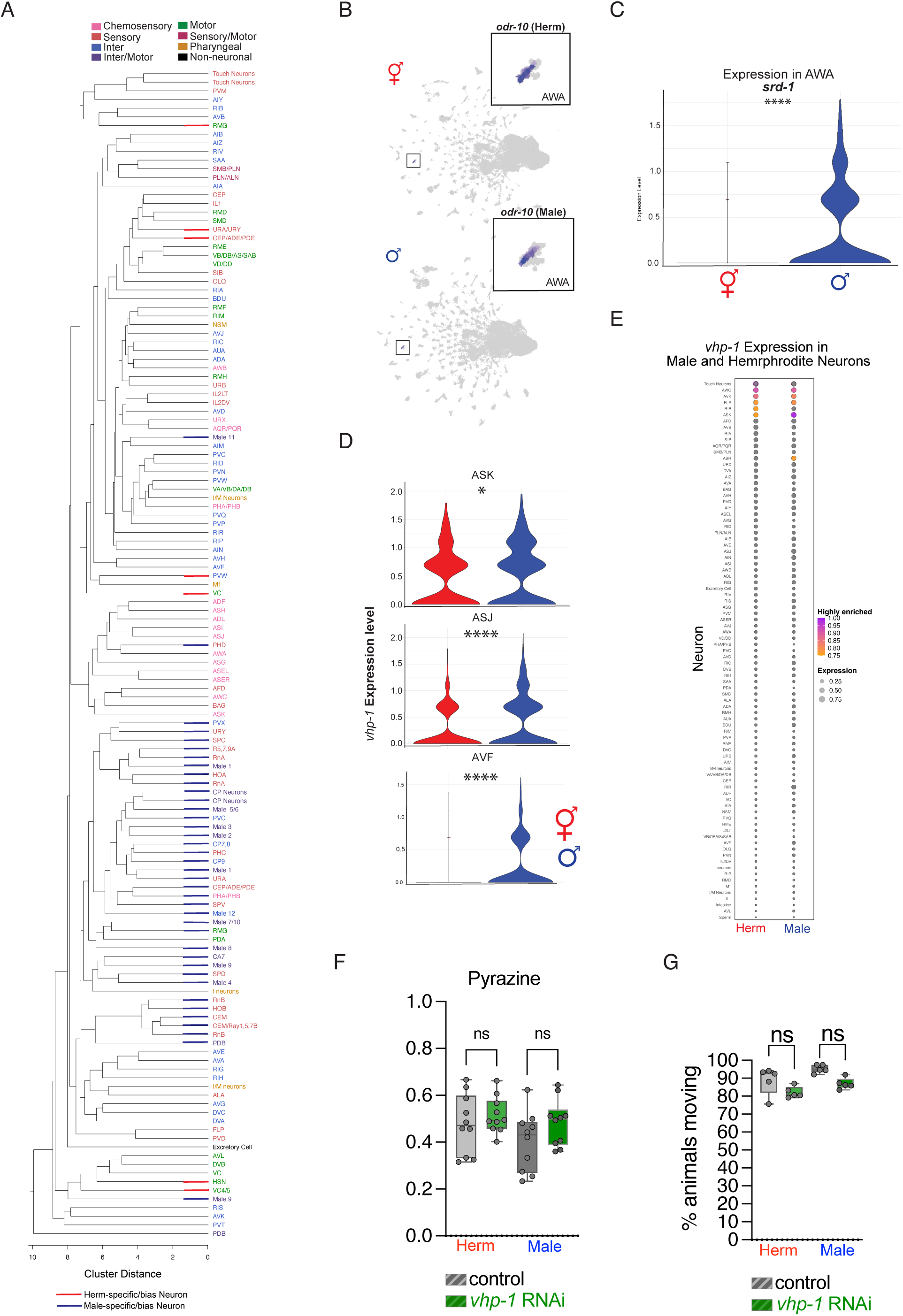
Sex-specific expression patterns and functional analysis of *vhp-1*. **(A)** Hierarchical dendrogram based on gene expression in each day 1 adult male and hermaphrodite neuron. Neurons are color-coded by functional subtype. Blue bars on branches indicate male-specific/bias neurons. Red bars indicate hermaphrodite-specific/bias neurons **(B)** Feature plot of *odr-10* expression in both sexes highlights enrichment in AWA and higher levels in hermaphrodites **(C)** Expression level of *srd-1* in AWA, showing upregulation in males. **(D)** *vhp-1* is upregulated in males in the AVF, ASK, and ASJ neurons. Adjusted p-values from Wilcoxon Rank Sum test. *p<0.05, ****p<0.0001. **(E)** Average normalized expression of *vhp-1* across neurons by sex. Dot size represents normalized expression values. Neurons with highest expression (≥ 0.75) are highlighted in orange (minimum) and purple (maximum). For reference the mean expression across all genes is ∼0.02 **(F)** Neuron-specific RNAi knockdown of *vhp-1* does not affect pyrazine chemotaxis in either sex compared to vector control or **(G)** Animals locomotion compared to vector control. One-way ANOVA with Bonferroni post hoc analysis ****p<0.0001, ns: p > 0.05.

## Notes

### Competing Interest Statement

The authors have declared no competing interest.

### Summary of Updates

Additional data, functional analyses, data analysis, and annotation refinements.

